# Alternative splicing and gene expression repurpose ancestral genes through distinct evolutionary routes

**DOI:** 10.64898/2026.09.05.749416

**Authors:** Federica Mantica, Antonio Torres-Mendez, Irene Bódalo-Zapata, Luis Pedro Iñiguez, Yamile Marquez, Violeta Morin, Jean-Yves Roignant, Manuel Irimia

**Author notes:** Co-corresponding authors: **Federica Mantica**, Centre for Genomic Regulation (CRG) Dr. Aiguader, 88, 08003 Barcelona, Spain; **Manuel Irimia**, Universitat Pompeu Fabra, Dr. Aiguader, 88, 08003 Barcelona, Spain. Neural Circuits & Evolution Laboratory, The Francis Crick Institute, London NW1 1AT, UK.

## Abstract

Specialized tissues often evolve by repurposing ancestral genes. Tissue-specific alternative splicing (TS-AS) and tissue-specific gene expression (TS-GE) have been proposed as complementary mechanisms contributing to tissue specialization, but whether they provide broadly interchangeable solutions or instead follow distinct, constrained evolutionary routes remains unknown. To address this question, here we compare TS-AS and TS-GE across eight homologous tissues, 20 bilaterian species and 7,178 ancestral gene families spanning 700 million years. The two mechanisms were similarly prevalent and, together, they affected nearly three-quarters of ancestral gene families across our dataset. However, they targeted largely non-overlapping gene sets within species, with contrasting architectures and functions. TS-AS usually acts on longer, exon-rich genes and modulates intracellular machinery shared across tissues, whereas TS-GE preferentially evolves in duplicated genes and impacts tissue-defining extracellular and nuclear processes. Unexpectedly, although TS-GE was globally more conserved, neural-specific splicing gains outnumbered expression gains at the origins of vertebrates and insects, revealing a prominent contribution of TS-AS to early nervous system transcriptome diversification. Exon-level orthology reconstructions further uncovered extensive recurrent evolution of neural-specific exons in the same ancestral genes across the phylogeny. Importantly, deletion of two such microexons in *Drosophila* negatively impacted overall fitness and led to neurological-associated phenotypes such as reduced climbing or hyperactivity. Altogether, our findings reveal distinct but complementary evolutionary routes by which gene expression and alternative splicing repurpose ancestral genes, and highlight recurrent neural exon evolution as a functionally important contributor to nervous systems.

## Introduction

Repurposing of ancestral genes for new biological roles is a pervasive process in animal evolution. Many tissue- and cell type-specific functions indeed arise through regulatory modification of conserved genes rather than through the emergence of novel ones ^1^. Two molecular mechanisms are key in mediating such gene repurposing: the gain of tissue-specific gene expression (TS-GE) and tissue-specific alternative splicing (TS-AS). These mechanisms have the potential to generate analogous functional outcomes resulting in specialization of tissue-specific proteomes. However, they achieve these outcomes in different ways ^2^. TS-GE gains are often acquired following gene duplication, where one gene paralog restricts its expression to a specific tissue ^3–5^. By contrast, TS-AS usually arises through the evolution of tissue-specific exons that generate alternative isoforms ^6^, with the exon-containing isoform being preferentially represented in a particular tissue. Thus, both TS-GE and TS-AS have the potential to specialize the transcriptome in one tissue while preserving the ancestral state in all others. However, whether this molecular analogy translates into equivalent and interchangeable evolutionary roles, or whether gene architecture and function constrain which regulatory route is favored, remains unclear.

Evidence for the evolutionary relevance of these mechanisms comes from both targeted and large-scale studies. Single-gene analyses have shown how TS-GE and TS-AS likely contributed to key innovations (e.g. ^7–9^). At a broader scale, multiple genome-wide analyses of TS-GE across ancestral mammalian, vertebrate, and bilaterian genes have linked gains of expression specificity in ancestral genes to the emergence of phenotypic features at the level of tissues and cell types ^4,5,10–16^. In contrast, genome-wide comparative studies of alternative splicing are scarcer and have generally covered shallower phylogenetic distances. Characterizations of alternative splicing landscapes across tissues have been carried out within tetrapods and mammals ^17–21^, as well as in a focused evolutionary comparison of neural TS-AS microexons between mouse and fruit fly ^22^. These studies have jointly revealed a much faster evolutionary turnover for alternative splicing than gene expression profiles across species and tissues. Nevertheless, whether this rapid turnover predominantly reflects species-specific TS-AS exons with limited functional relevance, or instead represents a substantial source of lasting evolutionary innovation has not been systematically investigated across deep evolutionary timescales.

Therefore, two important knowledge gaps remain. First, TS-AS has been comparatively understudied relative to TS-GE. This was in part due to technical challenges, including robust quantification of alternative splicing and the establishment of reliable exon orthologies across species. The latter is essential not only to reconstruct the evolution of tissue-specific exons, but also to distinguish true conservation from the independent, convergent evolution of different exons within homologous genes. As a result, the relative contributions of conservation, lineage-specific turnover and convergence to TS-AS evolution remain poorly understood. Second, direct comparisons between TS-AS and TS-GE are lacking, particularly across large evolutionary scales and within a unified analytical framework. This effectively hinders a comprehensive understanding of their relative contributions, evolutionary interplay and the intrinsic biases that favor one regulatory route over the other.

Here, we conducted the first systematic analysis of the evolution of TS-AS across deep phylogenetic distances, focusing on a balanced phylogeny for two major animal clades, vertebrates and insects, comprising a total of twenty bilaterian species (**Fig. 1a**, top; see **Supplementary Table 1**). For these species, we previously generated a bulk RNA-seq dataset spanning eight homologous tissue types (neural, muscle, testis, ovary, excretory, epidermis, digestive, and adipose tissues; **Fig. 1a**, center-top), which we used to characterize TS-GE genes included in 7,178 bilaterian-conserved gene orthogroups (**Fig. 1a**, center-bottom) ^4^. Within the same framework, we now defined and characterized TS-AS exons, enabling a direct comparison between ancestral genes evolving TS-AS and those evolving TS-GE (**Fig. 1a**, bottom). This comparison unveiled similarities and differences in the molecular, functional, and evolutionary features of TS-GE and TS-AS genes across tissues, species and evolutionary clades, showing that both mechanisms are complementary but not fully interchangeable. In addition, we found an unexpected deep evolutionary role for TS-AS in the transcriptomic diversification of vertebrate and insect brains, and identified extensive convergent evolution of neural-specific exons. Functionally testing two of these convergent microexons in Drosophila revealed pronounced behavioural and fitness consequences, highlighting their functional relevance.

**Fig. 1.**
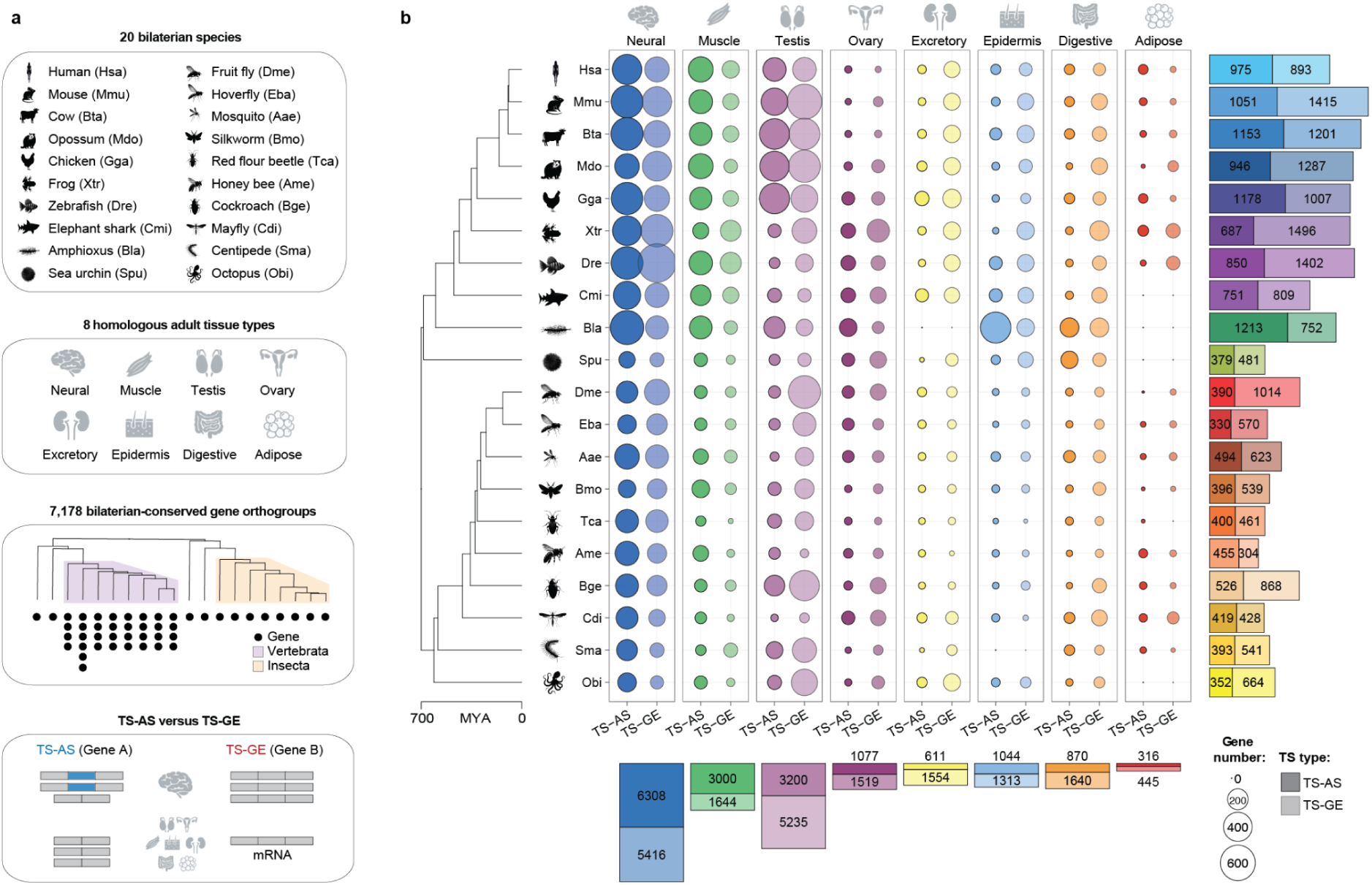
Framework overview and landscape of TS-AS and TS-GE genes: **a.** Schematic representation of the framework, showing the bilaterian species included (top), the adult tissue types analyzed (center-top), the gene orthogroups considered (with one generic orthogroup example depicted; center-bottom) and the regulatory mechanisms compared (bottom). **b.** Left: time-calibrated phylogenetic tree including the species scientific acronyms (see Methods for scientific names and **panel a** for common names), with evolutionary distances derived from TimeTree ^25^ (MYA: million years ago). Animal silhouettes were generated through *Bing Chat* by Microsoft (2023) [https://www.bing.com] and individually validated, while tissue icons were generated using *ChatGPT* by OpenAI (2026) [https://chatgpt.com]. Right: Dotplot and barplots representing numbers of TS-AS (solid shading) and TS-GE genes (transparent shading) across species (rows) and tissues (columns). Genes with double tissue-specificity (i.e., TS-AS or TS-GE in two tissues) are included twice in the dotplot but counted only once in the species-level barplot.

## Results

### Global landscapes of TS-AS and TS-GE genes across bilaterian animals

We first set out to compare global landscapes of TS-AS and TS-GE across bilaterian-conserved genes from the twenty species included in our phylogeny. We quantified alternative splicing levels using *vast-tools* ^23,24^ applied to the aforementioned multi-tissue, bulk RNA-seq dataset ^4^ (see **Supplementary Table 2**). We identified TS-AS events as those cassette exons in protein-coding regions showing tissue-specific inclusion (see **Methods** and **Extended Data Fig. 1a)**. TS-AS genes were defined based on the presence of at least one such exon, while TS-GE genes corresponded to the set with preferential tissue expression we had previously established ^4^ (**Supplementary Table 3**).

The numbers of TS-AS and TS-GE genes were remarkably similar within species (**Fig. 1b** and **Extended Data Fig. 1b**), but their phylogenetic distributions varied greatly. In general, chordates and vertebrates respectively exhibited higher numbers of TS-AS and TS-GE genes compared to other bilaterians (**Fig. 1b**), a pattern that remained solid after normalizing by the total number of ancestral protein-coding genes (**Extended Data Fig. 2a**). Relative TS-AS and TS-GE contributions also varied substantially across tissues. Neural samples exhibited the highest levels of both types of regulation (**Fig. 1b**, bottom), with a modest bias toward TS-AS in most species (**Extended Data Fig. 2b**). Muscle and testis also displayed substantial impact of tissue-specific regulation, but respectively skewed toward TS-AS and TS-GE (**Fig. 1b** and **Extended Data Fig. 2b)**.

### TS-AS and TS-GE genes are non-overlapping sets with differential features

We next directly compared TS-AS and TS-GE genes across all species and tissues. Remarkably, the two gene sets showed consistently low overlap within species (**Fig. 2a**). Although this pattern may be an underestimation influenced by the read coverage thresholds used to define TS-AS exons (see **Methods**), this minimal overlap nonetheless indicates that TS-AS and TS-GE may have largely complementary impacts on transcriptomes. Moreover, these gene sets systematically differed in intrinsic features that may be associated with, and potentially bias, the evolution of one mode of tissue-specific regulation over the other (see **Supplementary Dataset 1** for feature data).

**Fig. 2.**
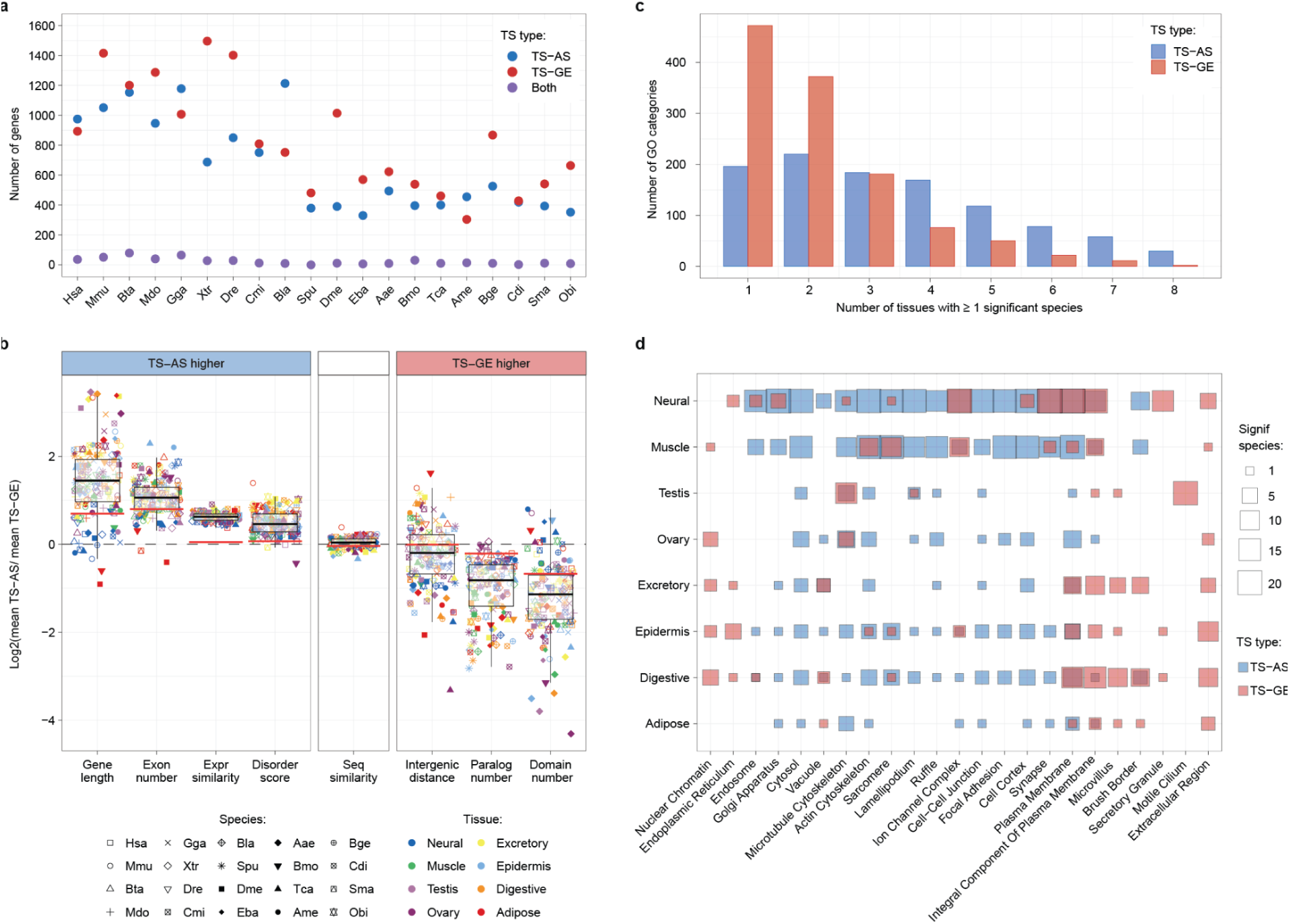
Differential features and functions of TS-AS and TS-GE genes: **a.** Number (y axis) of TS-AS (blue), TS-GE (red) and both TS-AS and TS-GE genes (purple) across species (x axis). **b.** Distributions of the log2 ratio between the average of each feature (x axis) in TS-AS and TS-GE genes in each species (shape) and tissue (color). The dashed, black line marks the baseline level (no differences between TS-AS and TS-GE genes) while the red bars represent the average of the medians of the same distributions obtained from 10 randomized TS-AS and TS-GE gene sets (see **Methods** and **Supplementary Fig. 1a**). **c.** Number of GO categories (y axis) based on the number of tissues (x axis) in which TS-AS (blue) and TS-GE (red) genes are enriched in at least one species. **d.** Number of species with significant enrichments for TS-AS (blue) and TS-GE (red) genes in each tissue (y axis) and selected cellular component GO categories (x axis; see **Supplementary Fig. 1b**).

First, TS-AS genes generally displayed more complex gene architecture than their TS-GE counterparts. They were consistently longer in all tissues except for neural (**Fig. 2b**), where TS-GE genes showed similarly large gene sizes in most species (**Extended Data Fig. 3a–h**). This is in line with the widely recognized tendency of neural genes to harbor particularly long introns ^26^. Importantly, however, TS-AS genes contained more exons than TS-GE genes in every tissue (**Fig. 2b**), including neural, revealing a robust association between exon-rich gene architecture and TS-AS propensity.

Second, beyond exon-intron architecture, these gene sets also differed in key evolutionary and structural features (**Fig. 2b** and **Extended Data Fig. 3a-h**). TS-AS genes exhibited more uniform expression profiles within gene families than TS-GE genes, and encoded proteins with (i) relatively more residues overlapping intrinsically disordered regions and (ii) fewer annotated protein domains, as expected based on ^27,28^.

Finally, TS-GE genes were characterized by longer intergenic distances as well as higher duplication levels, in line with the well-established associations between gene duplication and gains of tissue-specific gene expression profiles (see **Introduction**). Notably, despite these differences, TS-AS and TS-GE genes did not differ in overall sequence similarity to their orthologs and paralogs (**Fig. 2b**), although both groups were more diverged than other ancestral genes (**Extended Data Fig. 3a-h**).

Thus, TS-AS and TS-GE genes in all species and tissues universally differed in multiple features. Importantly, such differences are not recapitulated by randomized TS-AS and TS-GE gene sets (red line in **Fig. 2b, Supplementary Fig. 1a;** see **Methods**), supporting a genuine biological association between these properties and each type of tissue-specific regulation.

### TS-AS and TS-GE differentially shape cellular and molecular functions

We next assessed how both types of tissue-specificity impact cellular and molecular functions across species and tissues. We found that TS-GE genes were in general enriched for GO categories that were significant in only one or two tissues (**Fig. 2c**), while enriched categories among TS-AS genes frequently recurred across multiple tissues (60% in three or more tissues vs 29% for TS-GE, P-value < 2e-16, Fisher’s Exact test). Therefore, TS-GE seems to preferentially target non-overlapping functional programs across tissues, whereas TS-AS modulates a smaller set of processes that are reused and differentially tuned.

To investigate these differences more concretely, we then focused on GO enrichments for selected cellular component terms, ordered to roughly mirror the transition from the cell interior to the extracellular space (**Fig. 2d**, **Supplementary Fig. 1b** and **Supplementary Fig. 2**). This analysis revealed a clear and unexpected compartmentalization of regulatory modes. Nuclear chromatin and the endoplasmic reticulum were exclusively enriched among TS-GE genes, whereas intracellular compartments such as the cytosol, Golgi, and endosomal system showed prevalent and consistent enrichments among TS-AS genes. A sharp transition emerged at the cell periphery: intracellular membrane-associated structures, from the cytoskeleton to the cell cortex, were predominantly shaped by TS-AS regulation, while extracellular space and extracellular region were exclusively associated with TS-GE. The plasma membrane and related categories, including synapses and ion channels, represented intermediate cases, displaying strong enrichment for both regulatory layers.

We also observed similar patterns for molecular function and biological process GO terms (**Supplementary Figs. 3, 4**). TS-AS genes were again preferentially enriched for intracellular regulatory and structural functions. Examples included kinase activity, cytoskeletal organization, and regulation of GTPase activity and of actin polymerization, with the strongest signals displayed in neural and muscle. In contrast, TS-GE genes were primarily associated with tissue-defining functions, such as neurotransmitter transport in neural tissue and spermatogenesis in testis. Notably, the limited subset of terms that was shared across all tissues was related to DNA-binding transcription factor activity (**Supplementary Fig. 3**). Together, these results show that TS-GE and TS-AS preferentially operate on complementary functional layers, structured across tissues and cellular compartments: TS-GE shapes distinct programs within each tissue, whereas TS-AS tunes shared intracellular machinery.

### TS-AS and TS-GE are widespread across ancestral bilaterian gene families

We next aimed to quantify the potential of these mechanisms to shape tissue-specific transcriptomes across the phylogeny. Strikingly, by combining all orthogroups displaying either form of tissue-specificity, we found that nearly 75% of ancestral bilaterian gene families are affected by at least one of them in at least one species (lineplots in **Fig. 3a**), underscoring their remarkable evolutionary potential for tissue specialization. We then asked whether TS-AS and TS-GE tend to arise in the same orthogroups across evolution, or whether each mechanism preferentially affects distinct gene families, as suggested by the very low overlap observed within species (**Fig. 2a**). While 25% and 20% of orthogroups respectively contain only TS-AS or only TS-GE genes, 30% evolved both types of regulation, even if usually in different sets of species (**Fig. 3a,b**) and with a stronger relevance for testis-specific transcriptomes (**Extended Data Fig. 4a**). These three orthogroup classes were enriched for distinct sets of gene functions (**Fig. 3c,d** and **Supplementary Table 4**), as was the orthogroup class that did not show any tissue regulation across species, which was largely restricted to core housekeeping roles (**Fig. 3d**).

**Fig. 3.**
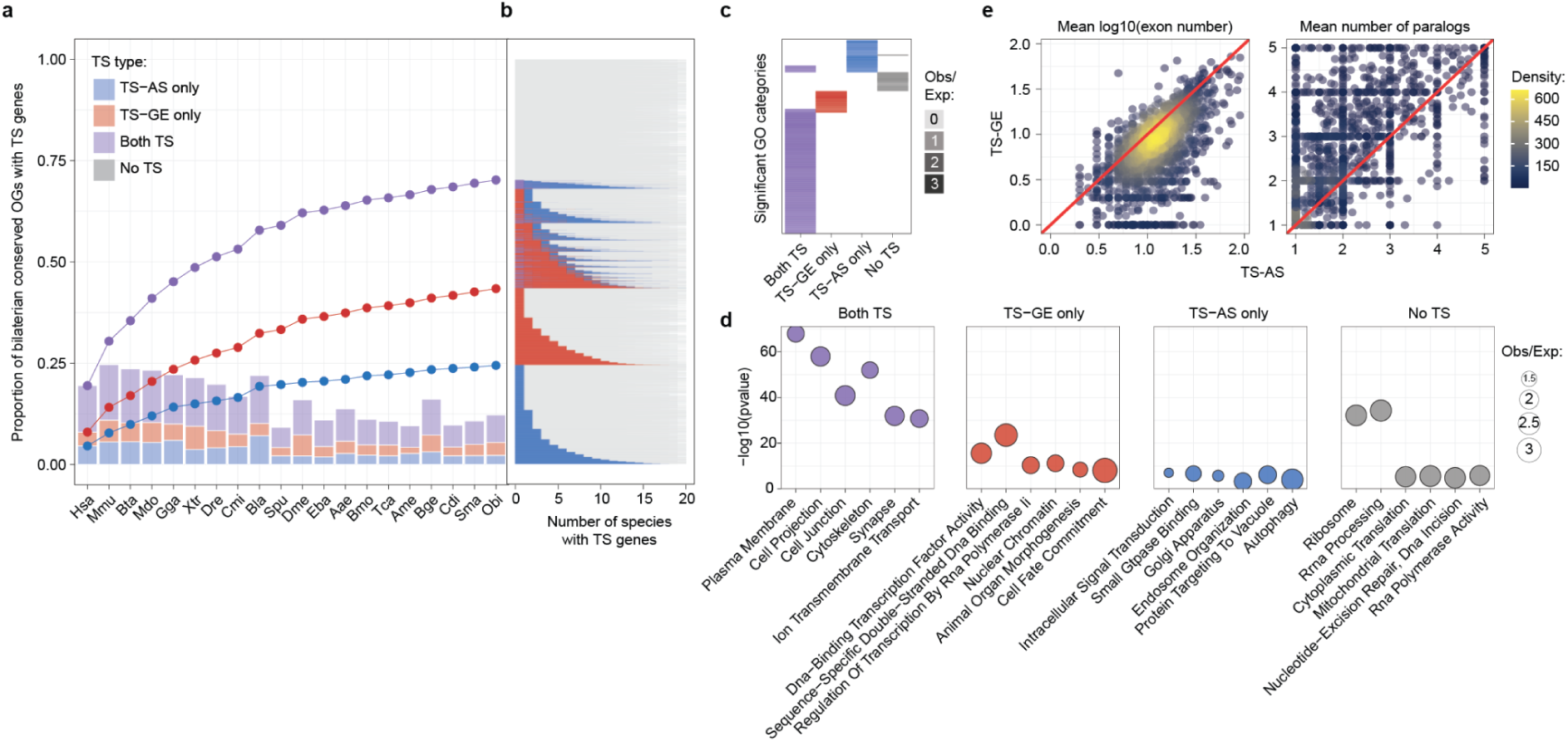
Characterization of TS-AS and TS-GE bilaterian-conserved gene orthogroups. **a.** Barplot representing the proportion of bilaterian-conserved gene orthogroups (y axis) including only TS-AS genes (blue), only TS-GE genes (red) or both TS-AS and TS-GE genes (purple) from each species (x axis). The line plots represent the stacked, cumulative distributions of these gene orthogroups categories across species. **b.** Distributions of the number of species including TS-AS (blue), TS-GE (red), both TS-AS and TS-GE (purple) or no TS genes (gray) across the gene orthogroups shown in panel a. **c.** Heatmap showing significantly enriched GO categories (hypergeometric test, P-value ≤ 0.001) across gene orthogroups containing only TS-AS, only TS-GE, both TS-AS and TS-GE, or no TS genes. **d.** -log10(P-values) for enriched selected GO categories from each orthogroup class shown in panel c (x axis). Point size indicates the observed-to-expected ratio, calculated as the proportion of GO-annotated genes in each group relative to background proportions (i.e., all gene orthogroups). **e.** Scatter plots of gene orthogroups containing both TS-AS and TS-GE genes, comparing mean log10 exon number (left) and mean number of paralogs (right) between TS-AS (x axis) and TS-GE (y axis) genes. Each point represents one orthogroup, and the red line marks the diagonal.

Notably, gene orthogroups with both types of tissue-specific regulation offered a unique opportunity to evaluate which features may predispose homologous genes to evolve TS-GE instead of TS-AS, or vice versa. In line with genome-wide associations (**Fig. 2b**), TS-AS genes had consistently more exons and were less duplicated than their TS-GE homologous counterparts (**Fig. 3e**). This suggests that differences in exon number and duplication events may bias genes toward distinct regulatory fates even within the same orthogroups.

In summary, TS-GE and TS-AS have broadly shaped ancestral bilaterian gene families through partially distinct evolutionary trajectories, associated with different gene features and functional profiles.

### Evolutionary conservation of TS-AS and TS-GE across bilaterians

To obtain initial insights into the relative evolutionary conservation of TS-AS and TS-GE, we compared orthogroup-level overlap of tissue-specific regulation within each tissue, both between the two regulatory mechanisms and between phylogenetic groups (see **Methods** and **Extended Data Fig. 4b**). First, shared regulation was higher for TS-GE than for TS-AS in all tissues (**Fig. 4a**, top; ∼2.5-fold greater median overlap across tissues; P-value ≤ 0.02 for all tissues, paired Wilcoxon tests). In other words, TS-GE genes were more likely than TS-AS genes to have orthologs showing the same tissue-specific regulation. Second, orthogroup-level overlap was more pronounced in vertebrates than insects for both regulatory layers (**Fig. 4a**, bottom; P-value ≤ 0.02 for all tissues, paired Wilcoxon tests). Third, across tissue types, neural consistently exhibited the highest regulatory overlap (median of up to 56% in vertebrates), with muscle always ranking second except for insect TS-GE (**Fig. 4b** and **Extended Data Fig. 4c**).

**Fig. 4.**
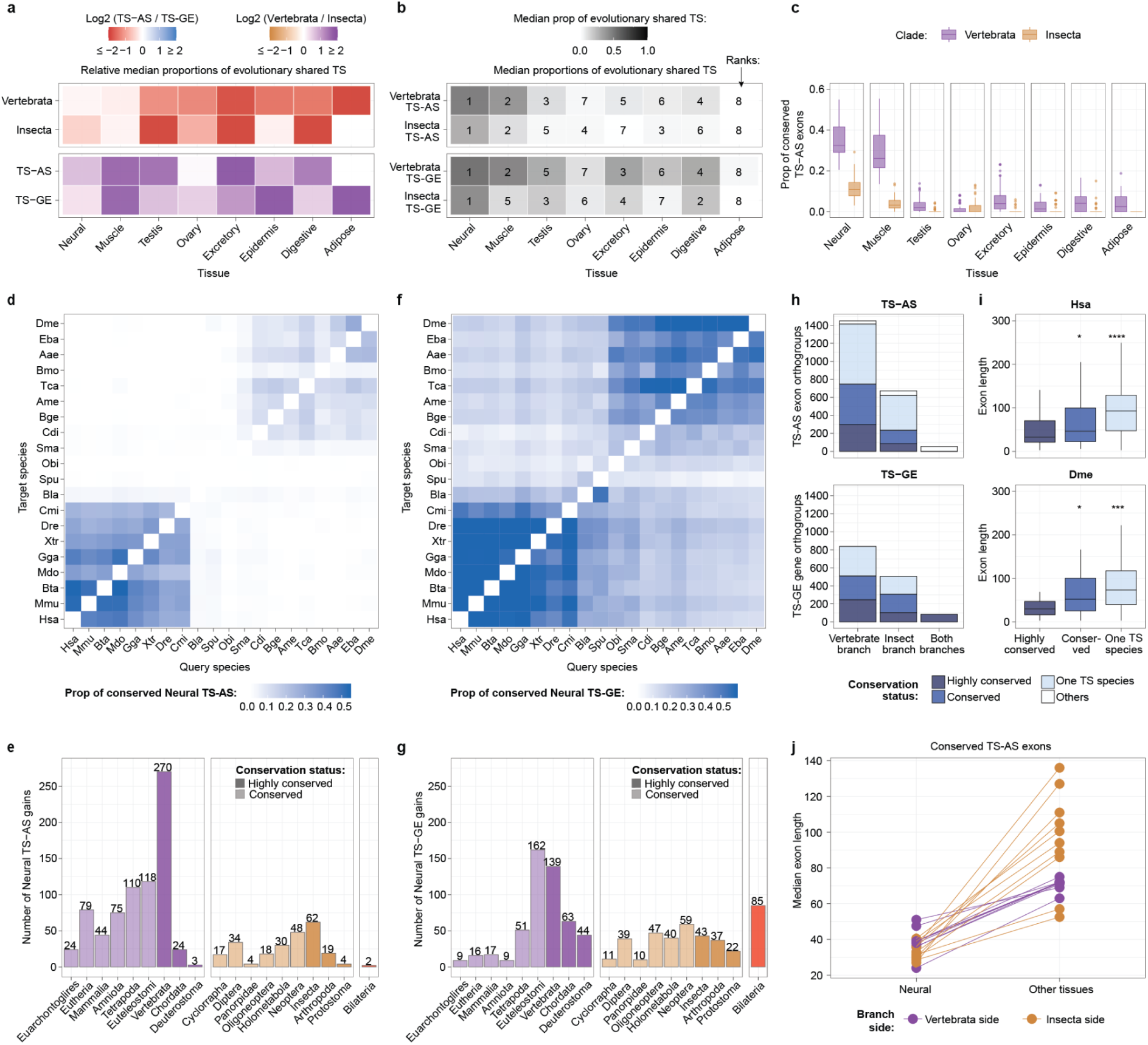
Evolutionary conservation of TS-AS and TS-GE across bilaterians. **a.** Patterns of shared tissue-specificity (i.e., TS overlap) within gene orthogroups. TS overlap was evaluated in a species-pairwise manner, and defined as the proportions of TS genes in the query species presenting at least one TS ortholog in the target species (see **Extended Data Fig. 4b,c**). TS overlap was computed both within regulatory layers and phylogenetic clades. Top: log2 ratio of median TS overlaps within regulatory layers (TS-AS/TS-GE) for each clade across tissues. Bottom: log2 ratio of median TS overlaps within phylogenetic clades (vertebrates/insects) for each regulatory layer across tissues. **b.** Median TS overlap in each tissue (x axis), stratified by regulatory layer (TS-AS vs TS-GE) and phylogenetic clade (vertebrates vs insects) (y axis). **c.** Proportions of conserved TS-AS exons (y axis) between all pairwise comparisons of vertebrates (purple) and insects (orange) across all tissues (x axis). **d,f.** Pairwise proportion of neural TS-AS exons (d) or TS-GE genes (f) in the query species (x axis) that are conserved in the target species (y axis) across all species combinations. **e,g.** Number of neural TS-AS (e) and TS-GE (g) gains across ancestral nodes, with conservation classes represented by different shades. **h.** Number of exon (top) or gene (bottom) orthogroups containing at least one neural TS-AS exon (top) or TS-GE gene (bottom) in each branch, with conservation classes represented by different shades. **i.** Distributions of exon lengths across conservation classes of exon orthogroups containing at least one neural TS-AS exon in human (top) and fruitfly (bottom). The significance levels plotted on top reflect the results of a *Wilcoxon* test comparing the highly conserved groups to each of the others, and correspond to the following P-value cutoffs: **** = P-value ≤ 0.0001; *** = P-value ≤ 0.001; * = P-value ≤ 0.05; ns = non significant. The y axis has been truncated for visualization purposes. Exon orthogroups classified as “Others” were excluded from this plot due to their low numbers. **j.** Median exon length of conserved TS-AS exons in neural (left) and all other tissues (right) across species. Points are colored by phylogenetic branch side, and paired values from the same species are connected by a line.

Importantly, however, while regulatory overlap within gene orthogroups is a proxy for regulatory conservation, this inference has a major limitation for TS-AS: true regulatory conservation can only be inferred when exon orthology is established, because different TS-AS exons may arise independently within orthologous genes. We thus ran *ExOrthist* ^29^ on our bilaterian-conserved genes to first infer exon orthogroups and then estimate TS-AS conservation, defined as the presence of same-tissue-specific orthologous exons within orthologous genes ^30,31^ (**Supplementary Datasets 2-3**). This allowed us to distinguish scenarios of true conservation from recurrent evolution.

In line with the higher shared gene orthogroup-level regulation (**Fig. 4b**), neural was by far the tissue with the highest TS-AS conservation, which was nearly negligible for the other tissues with the notable exception of muscle in vertebrates (**Fig. 4c, Extended Data Figs. 5, 6** and **Supplementary Figs. 5-9**). In particular, pairwise species comparisons revealed two prominent phylogenetic blocks of neural TS-AS conservation corresponding to vertebrates and insects (**Fig. 4d**). Remarkably, these blocks were reflected in an unexpectedly large number of exon orthogroups inferred to have acquired neural specificity in the vertebrate (n=270) and insect (n=62) ancestral nodes (**Fig. 4e** and **Supplementary Dataset 4**). A similar dual-block conservation pattern was observed for TS-GE (**Fig. 4f**), although with a stronger overall signal across the phylogeny, in line with the higher transcriptome-wide tissue conservation of gene expression vs. alternative splicing profiles (**Fig. 4a top,** and ^17^). Surprisingly, however, the number of genes inferred to have become neural TS-GE at the vertebrate (n=139) and insect (n=43) ancestral nodes was substantially lower than for TS-AS (**Fig. 4g**).

Despite these multiple ancestral gains, neural-specificity within exon orthogroups remained sparse: an average of 51.1% of neural exon orthogroups on each phylogenetic branch were restricted to a single species (**Fig. 4h** top; in contrast to 39.1% for neural TS-GE, **Fig. 4h**, bottom), and only two exons were confidently inferred to be bilaterian-ancestral (in contrast to 85 TS-GE neural genes, **Fig. 4f,g**). These exons corresponded to two microexons of respectively 6 and 9 nucleotides in *APBB1/2/3* and *MADD* (**Extended Data Fig. 7a,b**), two signal transduction proteins where such small in-frame insertions likely modulate protein-protein interactions ^32,33^. Moreover, neural exons with deep TS-AS conservation (i.e., since the vertebrate/insect ancestors or older) were significantly shorter than other neural exon groups (**Fig. 4i** and **Supplementary Fig. 10**). Similarly, conserved neural exons were systematically shorter than their conserved TS-AS counterparts in other tissues (**Fig. 4j**). All these patterns are consistent with the reported overall higher conservation of neural microexons in different species ^22,33,34^.

### Evolutionary recurrence of TS-AS and TS-GE across bilaterians

Beyond true TS-AS conservation, previous studies of alternative exon programs controlled by specific splicing regulators have revealed multiple cases of recurrent evolution, a scenario where distinct target exons independently emerged within orthologous genes ^22,29,35^. Motivated by these observations, we systematically assessed the extent of conservation vs. recurrent (or convergent) evolution of neural TS-AS in our dataset. We distinguished between cases where different neural exons evolved in the same protein-coding region at least twice across our phylogeny (type I convergence) and those independently arising at different positions within orthologous genes (type II convergence) (**Fig. 5a** and **Methods)**.

**Fig. 5.**
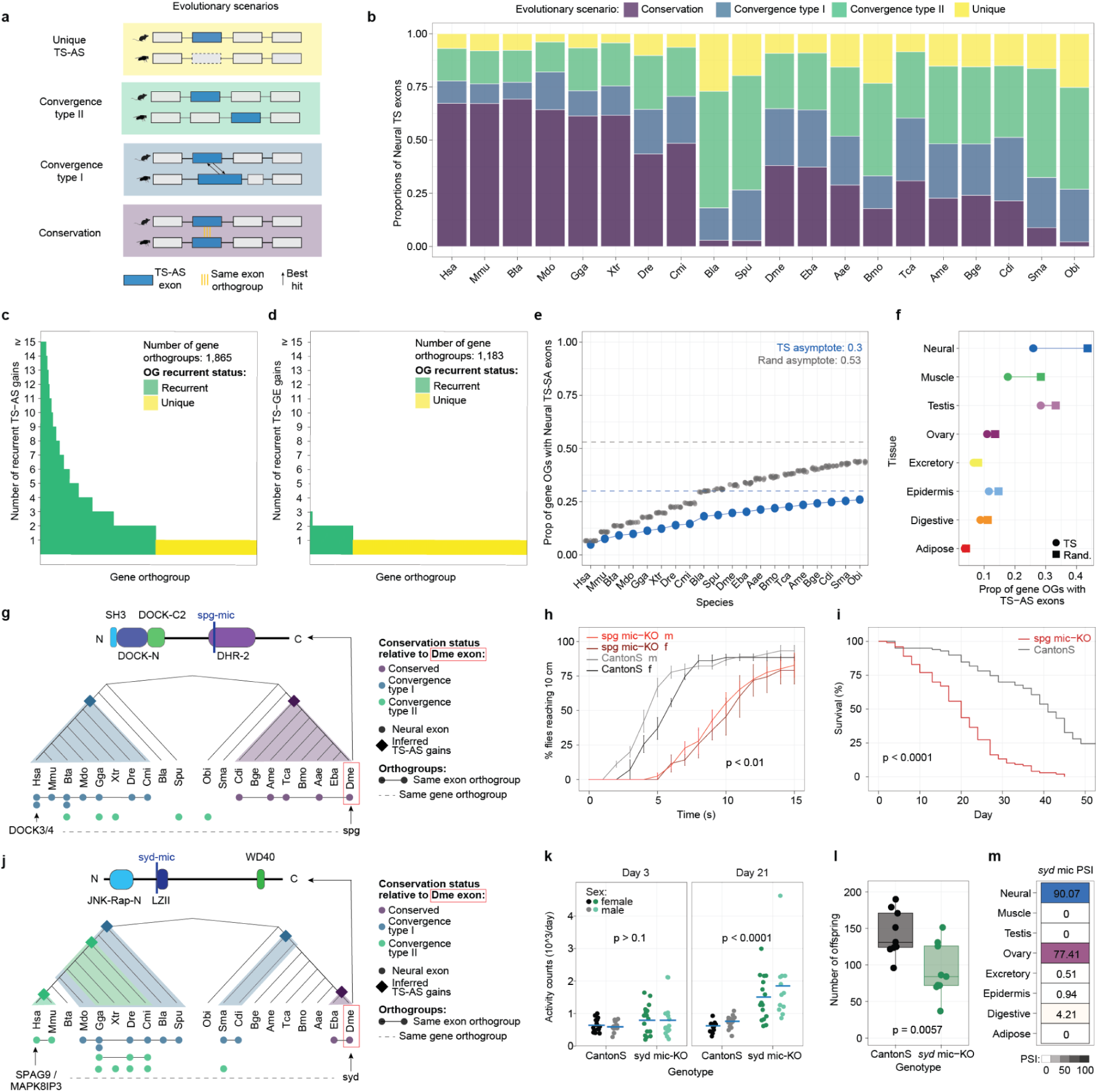
Evolutionary recurrence of TS-AS and TS-GE across bilaterians. **a.** Schematic representation of different TS-AS evolutionary scenarios. **b.** Proportions of neural TS-AS exons (y axis) across species (x axis) that are conserved, convergent type I, convergent type II or new. In case of different classification of the same exons between species pairs, priority is given to the highest inferred conservation category. **c,d.** Number of neural TS-AS (c) or TS-GE (d) gains within gene orthogroups containing at least one neural TS-AS exon (c) or TS-GE genes (d). Gene orthogroups are classified as “recurrent” when undergoing more than one TS gain. **e.** Cumulative distributions of the proportions of gene orthogroups (y axis) containing neural TS-AS exons either in the observed TS set (blue) or in 10 randomized sets (gray) across species (x axis). The horizontal dashed lines represent the asymptotes for the observed TS and randomized sets as predicted by nonlinear least-squares fitting of an asymptotic exponential model, y=*a*(1−e^−*b*x^), where *a* is the asymptote and *b* the rate constant. **f.** Proportions of gene orthogroups containing TS-AS exons in the observed TS set (circles) versus the median across 10 corresponding randomized sets (squares) for all tissues. **g.** Evolutionary history of TS-AS exons relative to the highlighted Drosophila microexon in spg. The position of the microexon within the encoded protein and its associated domain architecture are shown above. **h.** Anti-gravitaxis performance of 3-day-old CantonS control flies and spg microexon-KO flies. Points indicate the percentage of flies reaching 10 cm over time; error bars indicate the standard error of the mean. The P-value from a Mann-Whitney U test at 10 s is shown. **i.** Survival curves for 100 CantonS control flies and 100 spg microexon-KO flies. The P-value from a log-rank test is shown. **j**. Evolutionary history of TS-AS exons relative to the highlighted Drosophila microexon in syd. The position of the microexon within the encoded protein and its associated domain architecture are shown above. **k.** Locomotor activity of CantonS control flies and syd microexon-KO flies measured using a Drosophila Activity Monitor at days 3 and 21. Points represent individual flies and horizontal bars indicate group averages. P-values from Mann-Whitney U tests are shown. **l.** Number of offspring produced by age-matched female CantonS control flies and syd microexon-KO flies. Points represent individual females; the P-value from a Student’s t-test is shown. **m.** Percent-spliced-in (PSI) values of the syd microexon across tissues.

The relative proportions of conservation and convergence classes greatly vary across the phylogeny. In vertebrates, a median of 63% of neural TS-AS exons per species were conserved in at least another species, compared with 27% in insects and 3% in outgroup species (**Fig. 5b** and **Extended Data Fig. 8**). Strikingly, however, the vast majority of neural exons lacking direct conservation were not unique evolutionary events. Instead, they showed evidence of either type I or type II convergence with at least another species in our phylogeny (**Fig. 5b**). This implies that they arose in gene orthogroups that have also evolved neural-specific exons in other homologous genes, rather than representing a regulatory pattern that emerged only once in a given gene orthogroup.

This pattern was evident not only at the species level, but also across gene orthogroups. More than half of the orthogroups containing neural-specific exons presented at least one instance of convergent evolution across the phylogeny (**Fig. 5c**), in contrast with only 20% of gene orthogroups with neural TS-GE gains (**Fig. 5d**). Moreover, recurrent neural TS-AS was strongly concentrated in a restricted subset of ancestral genes rather than spreading randomly across the genome. Neural TS-AS accumulated in only ∼25% of bilaterian gene orthogroups and approached a predicted asymptote of ∼30%, substantially below randomized expectations (**Fig. 5e** and **Methods**). Altogether, this reveals strong constraints on the gene families prone to evolve neural TS-AS. Notably, such prominent convergent patterns and associated constraints were only observed in neural and, to some extent, in muscle tissues (**Fig. 5f, Extended Data Figs. 5, 6** and **Supplementary Figs. 5-9**).

Altogether, these results identify not only unexpectedly high levels of deep conservation of neural TS-AS in vertebrates and insects, but also a remarkable pattern of recurrent evolution of neural exons in orthologous genes across all bilaterians.

### Removal of recurrently evolved TS-AS exons in flies has functional consequences

These high levels of recurrent evolution of neural-specific exons suggest that TS-AS is of functional relevance for the targeted genes. To test this hypothesis we generated, as a proof of principle, exon-deficient fruitfly lines for two neural-specific microexons exhibiting different convergence patterns across our phylogeny, and performed a battery of assays to assess their overall fitness and behaviour (**Extended Data Fig. 9** and **Supplementary Table 5**).

The first one was an 18-nucleotide microexon located in the fruitfly *sponge* gene (*spg*). This microexon is conserved across insects and shows type I convergence in vertebrates (**Fig. 5g**). *spg* and its vertebrate orthologs *DOCK3* and *DOCK4* encode for guanine nucleotide exchange factors. They are involved in several neuronal processes such as cell migration, neuritogenesis and synaptic plasticity ^36–39^, and mutations in the human genes have been associated with neurological disorders and autism ^40–44^. The vertebrate and insect microexons convergently evolved within the catalytic DHR-2 domain ^45^ (**Fig. 5g**), and inclusion of the mouse counterpart has been shown to enhance the interaction with GRB2 ^46^. In line with these features, deletion of the *spg* microexon in fruitflies produced strong functional consequences *in vivo*. In negative gravitaxis assays, both male and female microexon-KO fruitflies showed markedly delayed climbing compared with control flies (**Fig. 5h**; P-value < 0.01, Mann-Whitney U test at time = 10 seconds), consistent with impaired locomotor or neurological function ^47^. Even more strikingly, loss of the microexon caused a profound reduction in lifespan, with median survival decreasing from 41 days in controls to only 20 days in knockout flies (**Fig. 5i**; log-rank test, P-value < 0.0001).

The second neural-specific exon was an 18-nucleotide microexon hosted by the fruitfly *sunday driver* gene *(syd),* which together with its vertebrate counterparts *SPAG9* and *MAPK8IP3* and orthologs in other lineages represented a hotspot of neural TS-AS evolution. In fact, the fruitfly microexon is conserved in the hoverfly, but independent, convergent events of either type I or type II have been inferred in the arthropod, deuterostome, vertebrate and human-mouse ancestors as well as in several single species (**Fig. 5j**). The fruit fly microexon is located immediately adjacent to leucine zipper II of *syd* (**Fig. 5j**), raising the possibility that its inclusion modulates interactions with kinesin-1, the dynein–dynactin complex and the small GTPase ARF6 ^48,49^, and thereby affects default neuronal transport or signalling ^50–54^. Consistent with a neural phenotype, *syd* microexon-KO flies developed a pronounced age-dependent hyperactivity that was absent in young adults and evident in both sexes by day 21 (**Fig. 5k**; P-value < 0.0001, Mann-Whitney U tests). In addition, microexon-KO flies produced significantly fewer offspring than age-matched controls (**Fig. 5l**; P-value = 0.0057, Student’s t-test), indicating a broader physiological impact that may include impaired fertility, potentially related to the lower but substantial inclusion of this microexon in ovaries (**Fig. 5m**).

In summary, these analyses support the idea that convergent evolution of TS-AS exons could provide an additional proxy for functional relevance, alongside direct evolutionary conservation.

## Discussion

In this work, we address two major gaps in the evolution of tissue-specific regulation. First, we characterized multi-tissue TS-AS landscapes across ∼700 million years of evolution, vastly extending the phylogenetic depth of previous comparative studies ^17–19^. Second, we directly compared TS-AS and TS-GE landscapes within a unified framework centered on ancestral bilaterian gene orthogroups, enabling us to assess their interplay within the same evolutionary context. This direct comparison is especially meaningful given the molecular and functional analogy between the two regulatory mechanisms. On one hand, TS-GE is typically associated with gene duplication and the generation of paralogous genes that can diverge in expression and function (often through specialization, see ^2^). On the other hand, the emergence of TS-AS exons effectively generates "internal paralogs", as distinct protein products are produced from a single gene without gene duplication. However, despite their similar molecular potential, our results show that TS-AS and TS-GE are in fact not interchangeable mechanisms for tissue specialization: they occupy largely different mechanistic, functional and evolutionary niches across bilaterians.

Mechanistically, it is not surprising that TS-AS genes exhibit significantly fewer paralogs compared to TS-GE genes. Unlike TS-GE genes, alternatively spliced genes have been shown to be less duplicated than the genomic background ^55–59^, despite some discordant studies ^60^. Importantly, these works generally quantified alternative splicing based on the presence or number of annotated isoforms. Here we instead specifically focus on highly tissue-specific exons, which likely strengthen the contrasting signal between alternatively spliced genes and background. Moreover, we showed that TS-AS genes tend to be consistently longer, exon-rich, and encode for more intrinsically disordered regions, suggesting that some of these features may reduce the likelihood of a successful duplication ^61,62^. More broadly, these findings also point to a general evolutionary model in which intrinsic gene architecture biases how tissue-specific regulation evolves: longer and structurally more complex genes may be more likely to diversify through TS-AS ^63,64^, whereas genes that are more amenable to duplication may preferentially acquire tissue specificity through TS-GE ^3–5^. This general model is also supported beyond bilaterian animals by previous analyses in *Arabidopsis thaliana*, where similar structural differences were observed between TS-AS and TS-GE genes ^65^.

Functionally, a key question concerns the relative impact of TS-AS and TS-GE across species and tissues. Perhaps unexpectedly, the overall number of genes affected by each of the two mechanisms is highly comparable within species. However, they represent largely non-overlapping sets with the distinct architectural properties described above. Consistently, their functional impacts also differ markedly. TS-GE primarily drives tissue-defining functions, often associated with extracellular or nuclear processes, whereas TS-AS preferentially modulates basic intracellular machinery that is shared across tissues. Interestingly, these differences are reflected in their tissue distributions. Neural stands out as the tissue most strongly shaped by TS-AS and TS-GE across the phylogeny, consistent with its reliance on both more tissue-defining synaptic and membrane functions and extensively regulated intracellular cytoskeletal and trafficking machinery. By contrast, muscle and testis respectively show a predominance of TS-AS and TS-GE, consistent with the prominent role of cytoskeletal and contractile functions in muscle and of tissue-restricted spermatogenic programs, including sperm flagellar machinery, in testis.

Evolutionarily, pairwise comparisons across tetrapod ^17^ or amniote ^18^ species, in which alternative splicing and expression values respectively drove species and tissue clustering, suggest that TS-AS is less evolutionarily conserved than TS-GE. Similarly, our analysis of shared regulation at the gene orthogroup level found a 2.5-fold higher overlap for TS-GE than TS-AS. This is at least in part because TS-GE includes many deeply conserved tissue-specific programs, some of which date back to the last bilaterian ancestor or earlier ^4^. However, this global pattern does not necessarily reflect the relative contribution of each mechanism at specific evolutionary transitions, during which TS-AS may still have played a comparatively greater role in lineage-specific innovation. This is particularly evident in the vertebrate ancestor, where we inferred a surprisingly high number of neural TS-AS gains, approximately twice as many as neural TS-GE gains (270 vs 139). This points to an unexpectedly large contribution for TS-AS in shaping the unique transcriptome of the emergent vertebrate brain, where microexons play a major part ^33^. In fact, small exons dominate the landscape of deeply conserved neural TS-AS in nearly all bilaterian species, and at least two cases of ancestral microexons conserved over 700 million years of evolution have been identified by our comparative analysis, despite the conservative criteria used to minimise false-positive ancestral assignments.

Moreover, the rapid evolutionary turnover of TS-AS does not necessarily imply that most lowly conserved exons represent isolated and weakly constrained innovations. Remarkably, we found that many neural-specific exons independently evolved in the same ancestral gene orthologs, either at equivalent protein-coding positions or elsewhere within the gene. These instances of convergent evolution affect a limited set of genes, indicating that the propensity to acquire neural TS-AS is likely constrained by protein function or protein architecture, as previously suggested for Tropomyosin ^66^ or FGFR-ligand interacting domains ^35^. Thus, although the specific exons may be evolutionarily flexible, the recurrent emergence of neural TS-AS within the same genes may reflect convergent solutions for modifying neural protein function in a similar manner. Such functional redundancy could, in turn, help explain their higher evolutionary turnover. While these hypotheses need to be formally tested through a systematic functional comparison of sets of convergent isoforms, the pronounced phenotypes caused by deleting two such recurrent neural exons in fruitfly are in line with this interpretation. Loss of a microexon in *spg* impaired climbing behaviour and markedly reduced lifespan, whereas deletion of a microexon in *syd* caused age-dependent hyperactivity and reduced offspring production. Therefore, these results provide a proof of principle that evolutionary convergence can help identify tissue-specific exons with important organismal functions.

Altogether, our findings support a model in which TS-AS and TS-GE have complementary but distinct roles in the evolution of tissue-specific transcriptomes and are not simply interchangeable. Their differential association with gene architecture, function, and conservation suggests that they contribute to tissue innovation through partly different evolutionary routes. Our observations provide key insights into how the vast majority of ancestral genes have been repeatedly repurposed into specialized functions throughout bilaterian evolution, and can pave the way for future back-to-back and large-scale evolutionary analyses of these two molecular mechanisms.

## Methods

### Genome and gene annotations

We focused on the twenty bilaterian species used in ^4^: human (*Homo sapiens, Hsa*), mouse (*Mus musculus, Mmu*), cow (*Bos taurus, Bta*), opossum (*Monodelphis domestica, Mdo*), chicken (*Gallus gallus, Gga*), tropical clawed frog (*Xenopus tropicalis, Xtr*), zebrafish (*Danio rerio, Dre*) and elephant shark (*Callorhinchus milii, Cmi*), amphioxus (*Branchiostoma lanceolatum, Bla*), sea urchin (*Strongylocentrotus purpuratus, Spu*), fruit fly (*Drosophila melanogaster, Dme*), marmalade hoverfly (*Episyrphus balteatus*, *Eba*), yellow fever mosquito (*Aedes aegypti, Aae*), domestic silk moth (*Bombyx mori, Bmo*), red flour beetle (*Tribolium castaneum, Tca*), honey bee (*Apis mellifera, Ame*), cockroach (*Blattella germanica*, *Bge*), mayfly (*Cloeon dipterum*, *Cdi*), centipede (*Strigamia maritima, Sma*), octopus (*Octopus bimaculoides, Obi*). The genome assemblies and gene annotation versions are reported in **Supplementary Table 1**. The gene annotations (GTFs) are available at https://data.mendeley.com/datasets/22m3dwhzk6/2.

### Bulk RNA-seq dataset

We used the bulk RNA-seq dataset released with ^4^, which comprises 1,222 samples spanning eight homologous tissues across the aforementioned twenty bilaterian species. From this dataset, we removed 6 samples in the honeybee (namely, the samples belonging to the honeybee Ovary_Forager_b and Ovary_Queen_d meta-samples) whose reads did not respect the minimum length requirements for AS quantification with *vast-tools* (see below). Metadata information for all samples is reported in **Supplementary Table 2**.

### Gene orthologies

We used the bilaterian-conserved gene orthologies generated in ^4^ by running *Broccoli v1.2* ^67^ on the reference proteomes (i.e., only including the isoform with the longest CDS sequence for each gene) of the 20 bilaterian species in our phylogeny. These include 7,178 gene orthogroups conserved in at least 12/20 species. These gene orthogroups are available at https://data.mendeley.com/datasets/22m3dwhzk6/2.

### Quantification of AS events using the *vast-tools* suite

Quantification of alternative splicing events was performed by running in succession different tools from the *vast-tools* suite (v2.5.1). We first ran *vast-tools align* on each of the 1,222 bulk RNA-seq samples in our dataset (see above). *vast-tools align* quantifies alternative splicing events using the PSI metric, defined as the proportion of mapping reads supporting exon inclusion ^68^ (see **Extended Data Fig. 1a**). For each species, we then ran *vast-tools merge* to integrate the PSI quantification across the samples belonging to the same meta-sample into a unique, representative value. This grouping procedure was performed to (i) increase read depth per meta-sample, (ii) dilute potential batch effects from publicly available samples, (iii) facilitate downstream analyses and comparisons by having a comparable number of replicates across tissues and species. For each species, we then ran *vast-tools combine* on the resulting meta-sample quantifications, generating one INCLUSION table. Finally, we further filtered these tables to only comprise CDS cassette exon events annotated within genes belonging to bilaterian-conserved orthogroups (see above).

### Definition of protein-coding TS-AS exons

We ran the *Get_Tissue_Specific_AS.pl* script from ^65^ on the INCLUSION tables described above, with the following parameters: *--min_dPSI 10 --min_dPSI_glob 25 --min_rep 1 -N 4 --event_type EX --test_tis “Neural,Testis,Ovary,Muscle,Kidney,Epithelial,DigestiveTract,Adipose”*. ***min_dPSI*** specifies the minimum delta PSI (dPSI) between the average PSI in the queried tissue and the average PSI in each of the other tissue groups; ***min_dPSI_glob*** specifies the minimum dPSI between the average PSI in the queried tissue and the average of the average PSIs in each of the other tissue groups. ***min_rep*** specifies the minimum number of replicates with coverage for a tissue group to be considered in the comparison. *-**N*** specifies the minimum number of tissue groups with coverage, that has to include the queried tissue. ***-test_tis*** specifies a comma-separated list of tissues for which a table with dPSI values (tissue - other tissues) will be printed. We selected the upregulated (UP) cassette exons defined as tissue-specific, and we applied further rules to more strictly associate them with one or more tissues. If the dPSI between the first and second tissues with the highest inclusion level was ≥ 15, the exon was considered tissue-specific in only the top tissue. Else, if the dPSI between the second and third tissue with the highest inclusion was ≥ 15, the exon was considered tissue-specific in the two top ranking tissues (double tissue-specificity). If none of the above conditions are respected, the exon is not considered tissue-specific even if it passes all requirements established by *Get_Tissue_Specific_AS.pl.* From the selected cassette exon events, we remove those where the tissue(s) with tissue-specificity include only VLOW or N samples as defined by *vast-tools* (i.e., very low coverage) ^23^. See **Extended Data Fig. 1a** for a schematic representation of the procedure.

### Definition of TS-AS and TS-GE gene sets

TS-AS genes in each species were defined as genes containing at least one TS-AS exon defined above. TS-GE genes in each species were defined as tissue-specific in ^4^, requiring a *Tau* ≥ 0.75, a minimum expression of 5 TPMs in at least one of the tissues and minimal expression differences between the tissue(s) with the tissue-specificity and all the others (see **Methods** and **Extended Data Fig. 2c** in ^4^). TS-AS genes, together with their defining TS-AS exons, and TS-GE genes are reported for all tissues and species in **Supplementary Table 3**.

### Generation of randomized TS-AS and TS-GE gene sets

The ten randomized TS-AS and TS-GE gene sets (**Fig. 2b** and **Supplementary Fig. 1a**) were generated as follows. For each species, tissue and randomization round, (i) TS-AS exons (from which TS-AS genes were defined) were randomly sampled from all exons passing the coverage thresholds set in *Get_Tissue_Specific_AS.pl* (see above), matching the size of the corresponding TS set; and (ii) TS-GE genes were randomly sampled from bilaterian-conserved genes passing the expression threshold used to define TS genes (≥ 5 TPM in at least one tissue), again matching the size of the corresponding TS set.

### Computation of gene features of TS-AS and TS-GE gene sets

The gene features shown in **Fig. 2b** and **Extended Data Fig. 3a-h** were computed as follows. Gene length, exon number and intergenic distance were derived from the reference GTF annotations (see above). Gene length (bp) was defined as the difference between the maximum end and minimum start genomic coordinates across all annotated entries for a gene, and exon number as the total number of annotated exons. Intergenic distance (bp) was defined as the sum of the strand-specific upstream and downstream genomic distances to the nearest neighboring genes on the same chromosome, computed using *bedtools closest ^69^*. The number of paralogs was defined as the count of genes from the same species assigned to the same gene orthogroup.

To compute the average sequence similarity and expression similarity, we used the pairwise similarity measures previously generated in ^70^, based on the same gene orthogroups (excluding Bla, Spu, Sma, and Obi) and RNA-seq dataset. These pairwise measures capture similarity at the level of protein sequence (sequence similarity) and expression profiles across tissues (expression similarities) between pairs of orthologous genes. For each gene, average sequence and expression similarity were calculated as the mean of all its pairwise similarity values with its orthologs.

Average disorder scores were computed by analyzing the reference protein sequences of the multi-exon genes of interest with *IUPred2A ^71,72^* using default parameters and averaging the resulting disorder scores at the amino-acid level. Finally, the functional domains of the multi-exon genes of interest were annotated using *pfam_scan.py v1.0* (https://github.com/aziele/pfam_scan, accessed on Oct 29th 2024) with *HMMER v3.3.2* (http://hmmer.org/) against the *Pfam-A* database (v32.0) ^73^, requiring an e-value ≤ 0.1. The number of functional domains was normalized by gene length before plotting (**Fig. 2b** and **Extended Data Fig. 3a-h).** All values for gene features across species are provided in **Supplementary Dataset 1**. The backgrounds shown in **Extended Data Fig. 3a-h** comprise all bilaterian conserved genes with the exclusion of TS-AS and TS-GE genes.

### GO enrichments of TS-AS and TS-GE gene sets

We performed Gene Ontology (GO) enrichment analyses using the GO annotations generated in ^4^, where we downloaded the human GO annotation (GeneID - GO mappings) from *Ensembl v106 ^74^* and supplemented it with the *ClueGO v2.5.5* level 5 annotations ^75^. To construct a human-based GO annotation file, we propagated GO terms from human genes to genes from the other species within the same orthogroup when at least 1/4 of the human genes in that orthogroup carried the corresponding GO annotation. Using these curated annotations, we performed GO enrichment analyses separately for each species (see **Fig. 2c,d and Supplementary Figs. 1b, 2-4**).

We tested enrichment of all TS-AS and TS-GE gene sets across all GO categories using a one-sided hypergeometric test implemented in R (*phyper(k − 1, K, N − K, n, lower.tail = FALSE)*). Here, *k* denotes the number of TS genes in the tested gene set annotated with the GO term, *K* the total number of genes annotated with the GO term, *n* the size of the tested gene set, and *N* the total number of bilaterian-conserved, protein-coding genes with any GO annotations in the species (background). We considered GO categories significantly enriched at P-value ≤ 0.001 and retained only categories annotated with at least 20 human genes for downstream analyses. Results from these analyses are shown in **Fig. 2c,d, Supplementary Fig. 1b** and **Supplementary Figs. 2-4.**

### Comparisons of gene orthogroups containing TS-AS and TS-GE genes

Gene orthogroups were divided into orthogroups containing only TS-AS genes, only TS-GE genes, both TS-AS and TS-GE genes or no TS genes. Gene lengths and paralog numbers were calculated as described above (see **Computation of gene features of TS-AS and TS-GE gene sets**). GO enrichment analyses were performed as described in the previous section, but collapsing gene-level GO annotations by gene orthogroup and using gene orthogroups, rather than individual genes, as the unit of analysis. All gene orthogroup-level GO enrichments are reported in **Supplementary Table 4** separately for each gene orthogroup class.

### Definition of exon orthogroups

For the gene orthogroups defined above, we ran *ExOrthist v1.2* ^29^ main module by (i) integrating all non-annotated exons from the vast-tools reference files with the *--extraexons* flag, (ii) adding all reciprocal *liftOver* hits (for the species pairs where these could be computed from available UCSC *liftOver* files) with the *--bonafidepairs* flag, and (iii) applying the default evolutionary conservation cutoffs for all evolutionary distances (short, medium, long). We then selected the resulting exon orthogroups (*EX_clusters.tab*) and performed an additional round of exon re-clustering, following the same logic as the original pipeline, to refine orthogroups containing at least one exon with a membership score ≤ 0.15. The resulting exon orthogroups (*EX_reclustered.tab*) were used for all downstream analyses after retaining only those belonging to bilaterian-conserved gene orthogroups and containing at least one TS-AS exon. These exon orthogroups are available in **Supplementary Dataset 2.**

### Definition of pairwise TS-AS and TS-GE conservation

A query TS-AS exon was considered conserved in a target species when presenting at least one TS-AS exon ortholog (with the same tissue-specificity) in the same exon orthogroup. Pairwise TS-AS conservation in each tissue (**Fig. 4d, Extended Data Figs. 5a, 6a, Supplementary Figs. 5a, 6a, 7a, 8a, 9a**) was assessed using the output of all pairwise species runs of *ExOrthist compare_exon_sets.pl* (*reg_exon_orth* category from the Compare_exons_summary_input-$query_species-$target_species.txt files), where the TS-AS exons in the species involved were given as an input. The proportion of conserved TS-AS exons was calculated relative to the total number of TS-AS exons in the query species. These files and other outputs of *ExOrthist compare_exon_sets.pl* are available in **Supplementary Dataset 3.**

Species pairwise conservation of TS-GE genes was computed starting from the inferred gains and losses of tissue-specificity (see ^4^). A query TS-GE was considered conserved in a target species if (i) both the query and the target species descended from the ancestral node where a tissue-specificity gain had been inferred and (ii) none of the species descended from an ancestral node where a secondary tissue-specificity loss had been identified.

### Inference of clade-specific gains of TS-AS and TS-GE regulation

For each tissue, we separately inferred the gain of clade-specific TS-AS regulation (**Fig. 4e, Extended Data Figs. 5c, 6c** and **Supplementary Figs. 5c, 6c, 7c, 8c, 9c**). For this purpose, exons were considered TS-AS both if they were directly classified as TS-AS and if they showed a strong tissue bias (*Tau* ≥ 0.6) with the highest exon inclusion occurring in the tested tissue. We inferred a gain of TS-AS regulation in a particular ancestor when in an exon orthogroup (i) TS-AS exons were present in the earliest diverging lineage(s) of the clade and in at least 50% of the clade and, (ii) tissue-specificity was absent outside the clade. To infer ancestral bilaterian tissue-specific exon orthogroups, we required the orthogroup to contain at least one TS-AS exon in vertebrates, one in insects, and one in each of the two outgroup lineages (amphioxus or sea urchin; centipede or octopus), with a minimum of nine TS-AS exons overall. These deliberately conservative criteria were chosen to minimise false-positive ancestral assignments, at the likely cost of some false negatives (as we address in the **Discussion**).

Several exon orthogroups displayed phylogenetic distributions inconsistent with the inferred ancestral states. To distinguish potential losses of TS-AS from independent gains in these cases, we compared exon lengths across species. We reasoned that small length differences preserving the open reading frame are more likely to reflect modifications of ancestral exons than independent origins. Exon orthogroups in which all TS-AS exons differed only by small length shifts that were multiples of 3 nt (up to 6 nt difference between the shortest and longest) were therefore interpreted as derived from the same ancestral exon and inferred to have gained tissue-specificity in the last common ancestor of all species containing TS-AS exons. The remaining exon orthogroups were classified either as single-species TS-AS (when only one species in the orthogroup contained TS-AS exons) or as bilaterian-shared but unresolved (“others”). See **Supplementary Dataset 4** for classification of TS-AS gains. In downstream analyses, exon orthogroups with inferred TS-AS gains in Vertebrata, Insecta, or older ancestors were labelled highly conserved, whereas orthogroups with more recent gains were labelled simply as conserved.

In case of TS-GE regulation, we directly used the tissue-specificity gains inferred in ^4^, from which we removed the species with inferred losses.

### Comparison of exon lengths across TS-AS-conservation classes

To test whether regulatory conservation was associated with exon architecture, exon lengths were compared across TS-AS conservation categories. Distributions of TS-AS exons in human and fruitfly across conservation categories in all tissues are reported in **Fig. 4i, Extended Data Figs. 5f, 6f and Supplementary Figs. 5f, 6f, 7f, 8f, 9f**, whereas distributions of neural TS-AS exons in all species are reported in **Supplementary Fig. 10**.

### Classification of TS-AS and TS-GE recurrence

To estimate the impact of recurrent evolution of TS-AS regulation in our dataset, each TS-AS exon was assigned to one of four evolutionary categories (**Fig. 5a**). <u>Conservation</u>: exons belonging to highly conserved or conserved exon orthogroups (see above). <u>Convergence type I</u>: exons occupying equivalent intronic positions in at least two species (corresponding to the BEST_HIT* entries in *ExOrthist*’s *Compare_exon_sets.pl* output) but not included in any exon orthogroup containing another TS-AS exon. <u>Convergence type II</u>: exons arising at non-orthologous intronic positions within orthologous genes. New: exons present in gene orthogroups where no other TS-AS exons are detected. When deriving species-wise classifications within gene orthogroups (**Fig. 5b, Extended Data Figs. 5g, 6g and Supplementary Figs. 5g, 6g, 7g, 8g, 9g**) evolutionary categories were assigned according to the highest-priority TS-AS exon present (priority order: conservation > convergence type I > convergence type II > new). For TS-GE, recurrent evolution was identified when more than one TS-GE gain had been inferred within a gene orthogroup.

### Saturation of TS-AS signal within gene orthogroups

The saturation of the TS-AS signal (**Fig. 5e** and **Extended Data Figs. 5j, 6j**) and the overall TS-AS signal within gene orthogroups was predicted by nonlinear least-squares fitting of an asymptotic exponential model, y=*a*(1−e^−*b*x^), where *a* is the asymptote and *b* the rate constant.

### Generation of CRISPR knockout (KO) lines

Mutant alleles for *spg* and *syd* were generated using the CRISPR/Cas9 system following the previously described procedure ^76^. Two independent guide RNAs (gRNAs) for each gene were designed using the gRNA design tool: http://crisprflydesign.org/. Oligonucleotides were annealed and cloned into pBFv-U6.2 vector (National Institute of Genetics, Japan). Vectors were injected into embryos of *y1 v1 P(nos-phiC31/int.NLS)X; attp40*. Transgenic flies were further crossed with *y2 cho2 v1 ; attp40 (nos-Cas9)/Cyo*, and flies from the F1 generation were PCR screened for the expected mutation using primer sequences flanking the gRNA sequences. *spg* allele was obtained using gRNA sequences (GCAATCGCCATCCATAAAAA and GATATACAACAAATGAACTAG) and produced a deletion of 110 nt comprising the microexon. *syd* allele was obtained using gRNA sequences (GTGTGTTTAGTAATTGCGAG and GGGTAATGAACTAATTCTAC) and produced a deletion of 155 nt comprising the microexon. Microexon-specific KO lines were isogenized with CantonS flies, which were used as controls for experiments.

### Phenotypic characterization of fruitfly microexon-KO lines

#### Longevity assay

The lifespan of 50 male and 50 female flies for each genotype was monitored every two days, and flies transferred to a new food vial every time. Experiments were done at 25°C. Flies that escaped were right-censored. P-value for the survival was calculated using the log-rank test.

#### Anti-gravitaxis assay

We separated 3-day-old males and females in three groups of 15 each under anesthesia one day before the experiment was performed. We transferred the flies to dry 25 ml glass graduated cylinders marked 10 cm from the bottom. We video recorded for 20 seconds after tapping the cylinders. We manually counted the number of flies crossing the mark for each second from the recorded videos. We repeated the experiments 3 times independently for the controls and mutant flies and averaged the data. We calculated P-values using Wilcoxon tests for the number of flies crossing the mark at 10 seconds.

#### Monitoring of locomotor activity and sleep patterns

We monitored male and female lines using Trikinetics *Drosophila* Activity Monitors (DAM) during 1-minute bins. Each fly was placed into a glass tube containing 2% agarose and 5% sucrose food. Flies were entrained for 4 days in 12:12 Light:Dark cycles (LD). All the experiments were performed at 25 °C. The sleep and activity parameters were analyzed using the MATLAB script SCAMP (https://academics.skidmore.edu/blogs/cvecsey/files/2019/03/Vecsey-Sleep-and-Circadian-Analysis-MATLAB-Program-SCAMP-2019_v2.zip)

#### Fertility assay

15 female and 15 male flies were placed on food vials and transferred to a new vial every three days. The total number of adult flies hatched was monitored daily for up to nine days. Experiments were done in triplicate at 25 °C. Student’s t-test was used for testing differences in the total progeny size between genotypes.

See **Supplementary Table 5** for raw data of all fly experiments.

## Data and code availability

The code used for all analyses is available on *GitHub* at https://github.com/fedemantica/bilaterian_AS. The Supplementary Datasets are available in Mendeley Data at https://data.mendeley.com/datasets/795wtjbyk9/1 (DOI: 10.17632/795wtjbyk9.1).

## Authors’ contributions

FM performed most computational analyses and generated all figures and tables, with the feedback of MI and technical contributions from LPI and IB-Z. AT-M performed all fly experiments. VM and J-YR generated the microexon KO fly lines. YM contributed with ideas and guidance. FM and MI wrote the manuscript with input from all other authors.

## Extended Data Figures

**Extended Data Fig. 1.**
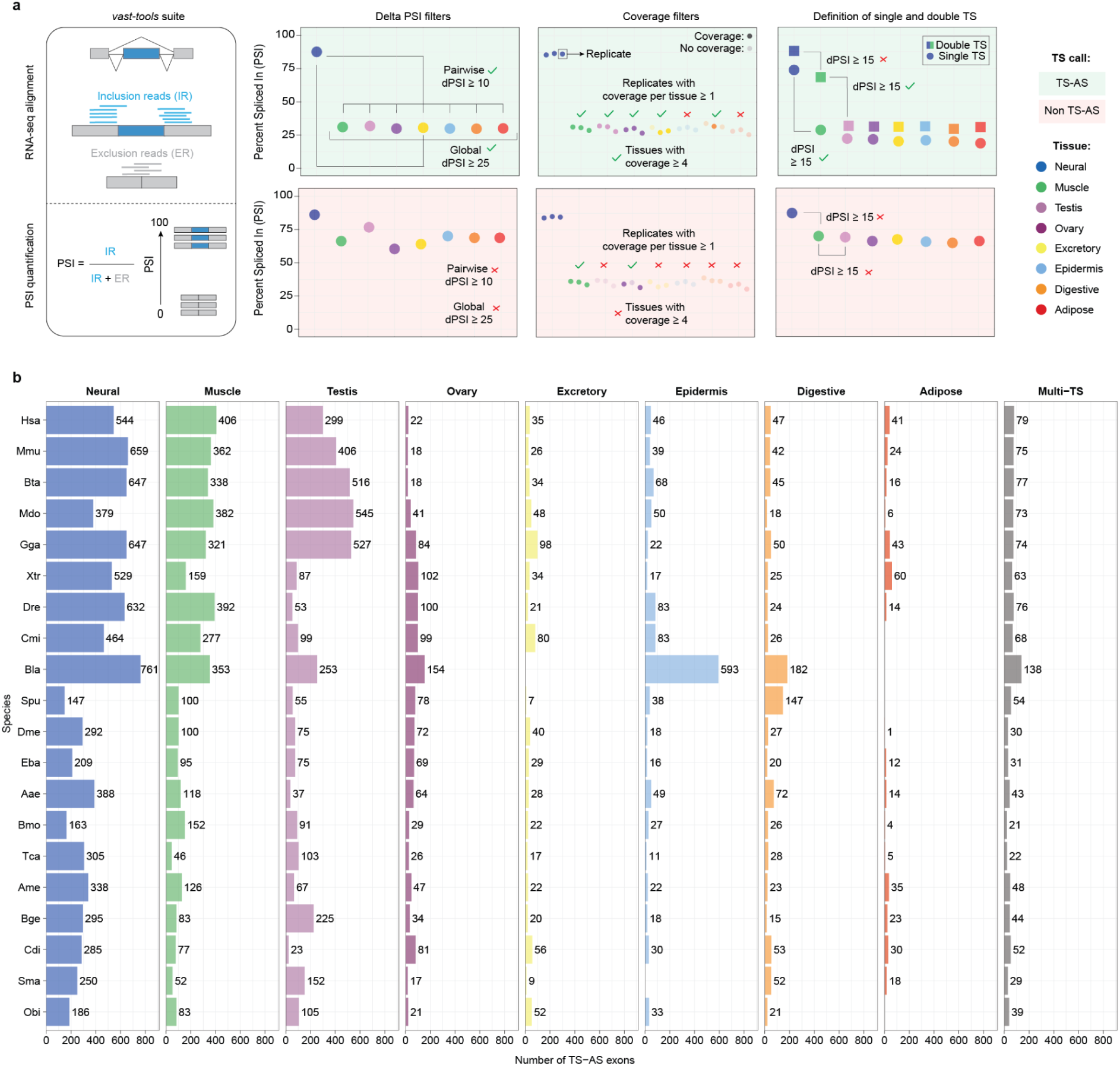
Definition of TS-AS exons: **a.** Schematic representation of the method adopted for the definition of TS-AS exons. **b.** Number of TS-AS exons across species (rows) and tissues (columns). Multi-TS indicate TS-AS associated with two different tissues (see panel a), and are also represented in the counts of the single tissues.

**Extended Data Fig. 2.**
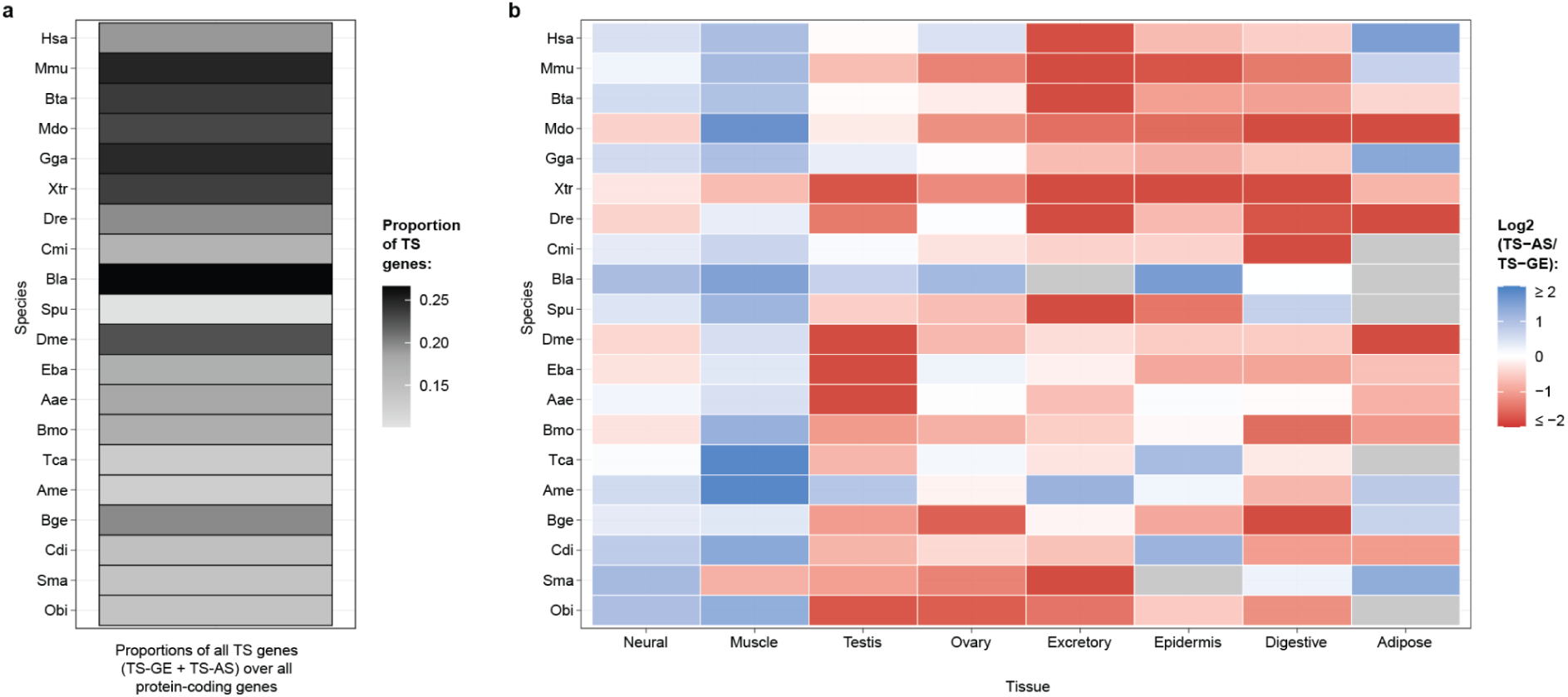
Relative proportions of TS-AS and TS-GE genes across species and tissues: **a.** Proportion of TS-AS and TS-GE genes over all ancestral protein-coding genes across species (y axis). **b.** Log2 ratio of the number of TS-AS to TS-GE genes across species (y axis) and tissues (x axis).

**Extended Data Fig. 3.**
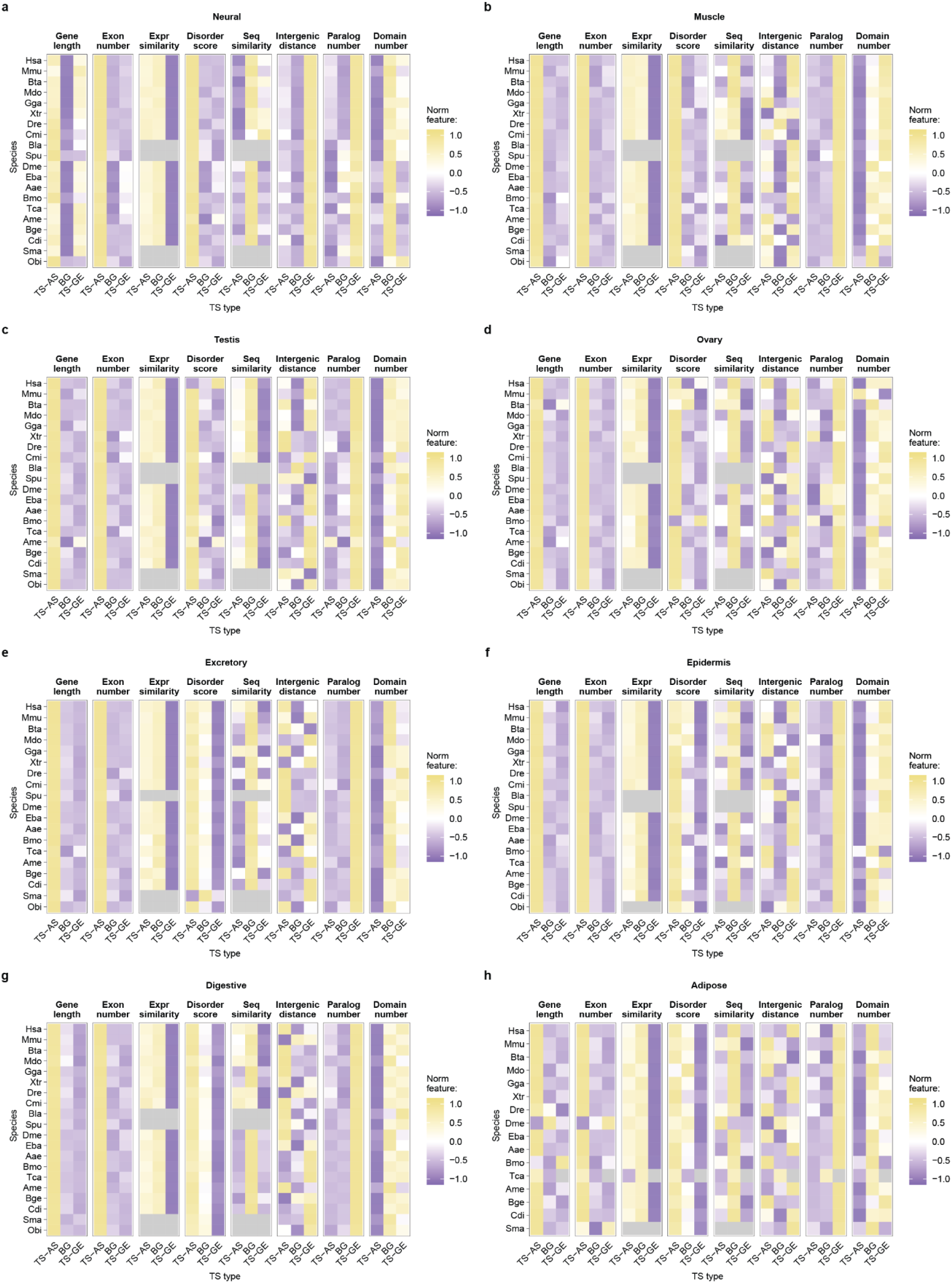
Features of TS-AS and TS-GE genes versus background. **a-h:** Z-scored features across TS-AS, TS-GE and background genes in each species and tissue. Background genes include all bilaterian ancestral protein-coding genes in each species minus TS-AS and TS-GE genes. See **Methods** for definition of all features.

**Extended Data Fig. 4.**
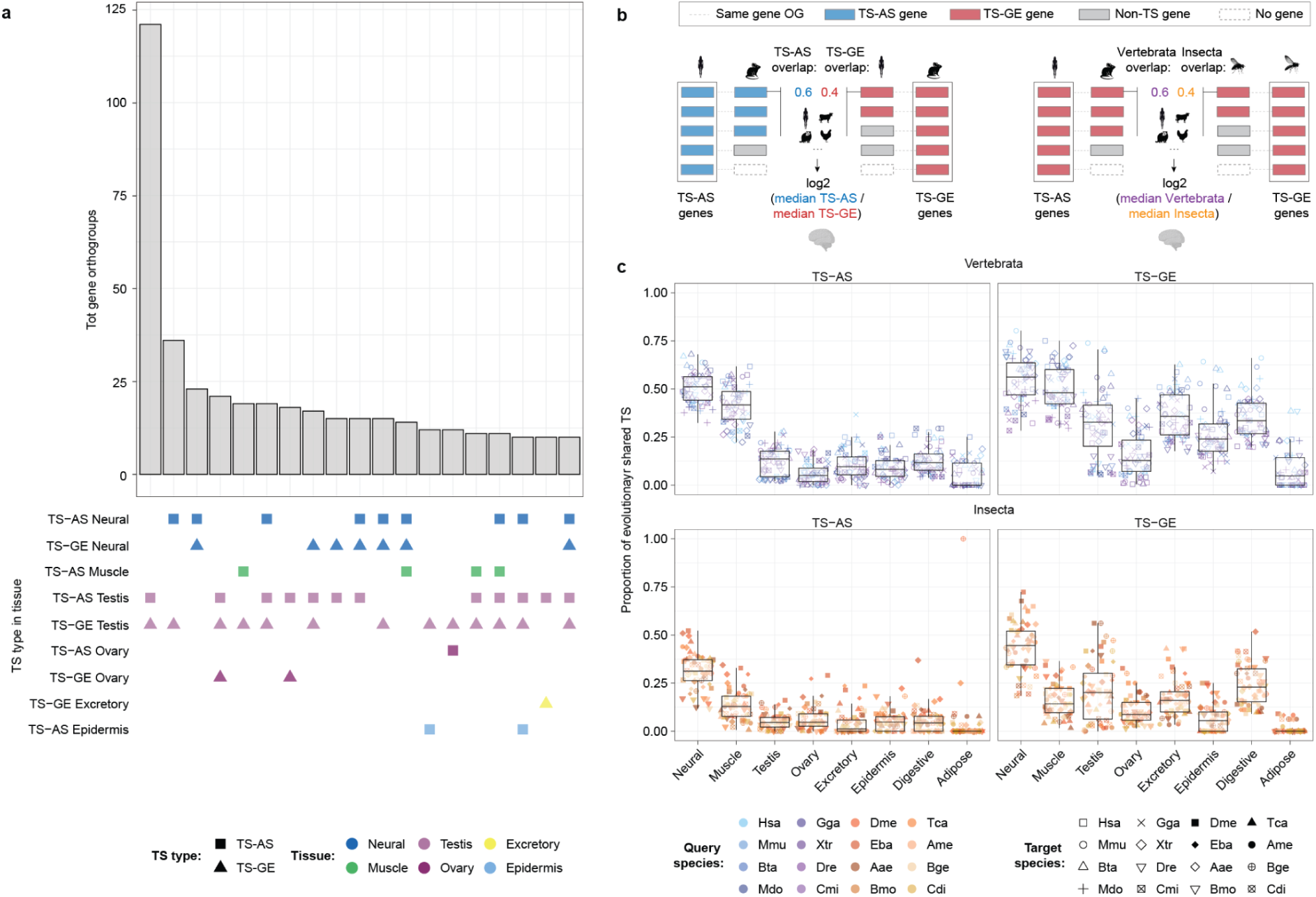
Co-occurrence of TS-AS and TS-GE genes within gene orthogroups. **a.** UpSet plot representing the ten most frequent combinations of TS types (shape) and tissues (color) within gene orthogroups containing both TS-AS and TS-GE genes. **b.** Schematic representation of the computation of species pairwise TS overlap within each tissue. TS overlap is computed separately between regulatory layers and within clades (left) and between clades and within regulatory layers (right). **c**. Proportion of bilaterian-conserved gene orthogroups containing either TS-AS or TS-GE genes in the query species (color) that also contain a corresponding TS-AS or TS-GE gene in each target species (shape) shown across tissues (x axis). Intra-vertebrata (top) and intra-insecta (bottom) comparisons were considered separately.

**Extended Data Fig. 5.**
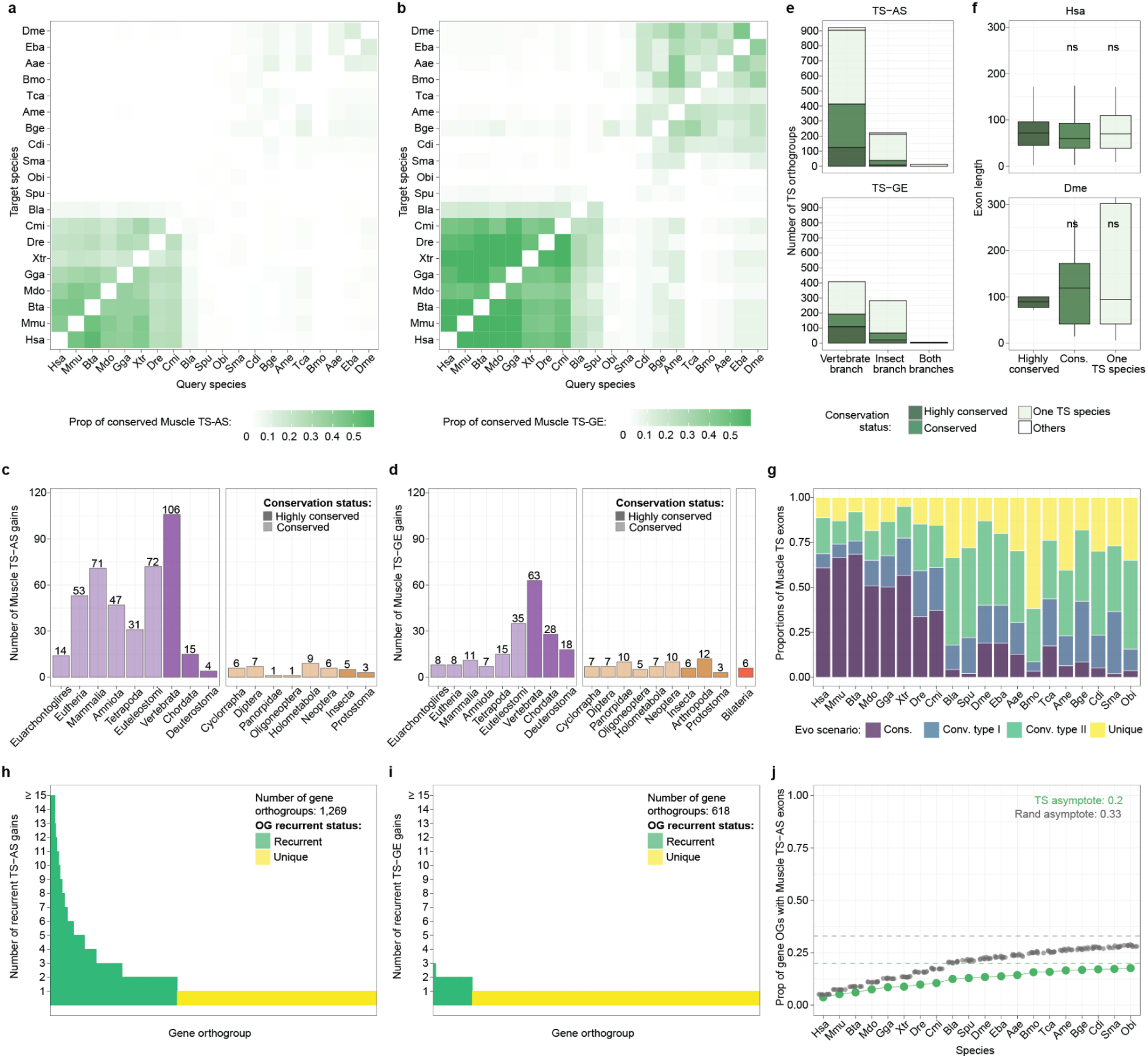
Evolutionary patterns of muscle TS-AS exons across bilaterians. **a,b.** Pairwise proportion of muscle TS-AS exons (a) or TS-GE genes (b) in the query species (x axis) that are conserved in the target species (y axis) across all species combinations. **c,d.** Number of muscle TS-AS (c) and TS-GE (d) gains across ancestral nodes, with conservation classes represented by different shades. **e.** Number of exon (top) or gene (bottom) orthogroups containing at least one muscle TS-AS exon (top) or TS-GE gene (bottom) in each branch, with conservation classes represented by different shades. **f.** Distributions of exon lengths across conservation classes of exon orthogroups containing at least one muscle TS-AS exon in human (top) and fruitfly (bottom). The significance levels plotted on top reflect the results of a *Wilcoxon* test comparing the highly conserved groups to each of the others, and correspond to the following P-value cutoffs: **** = P-value ≤ 0.0001; *** = P-value ≤ 0.001; * = P-value ≤ 0.05; ns = non significant. The y axis has been truncated for visualization purposes. Exon orthogroups classified as “Others” were excluded from this plot due to their low numbers. **g.** Proportions of muscle TS-AS exons (y axis) across species (x axis) that are conserved, convergent type I, convergent type II or new. In case of different classification of the same exons between species pairs, priority is given to the highest inferred conservation category. **h,i.** Number of muscle TS-AS (h) or TS-GE (i) gains within gene orthogroups containing at least one muscle TS-AS exon (h) or TS-GE gene (i). Gene orthogroups are classified as “recurrent” when undergoing more than one TS gain. **j.** Cumulative distributions of the proportions of gene orthogroups (y axis) containing muscle TS-AS exons either in the observed TS set (blue) or in 10 randomized sets (gray) across species (x axis). The horizontal dashed lines represent the asymptotes for the observed TS and randomized sets as predicted by nonlinear least-squares fitting of an asymptotic exponential model, y=*a*(1−e^−*b*x^), where *a* is the asymptote and *b* the rate constant.

**Extended Data Fig. 6.**
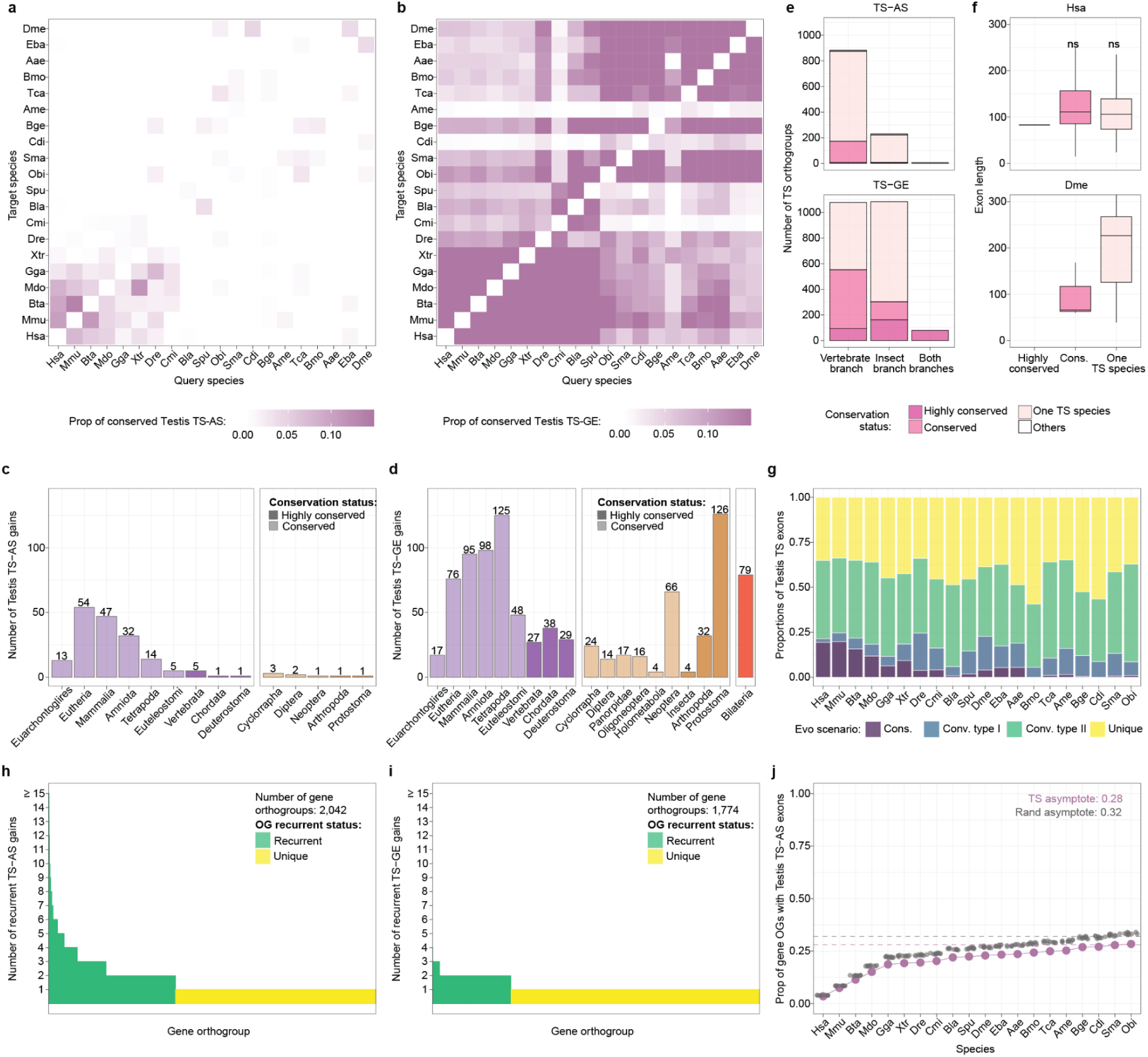
Evolutionary patterns of testis TS-AS exons across bilaterians. **a,b.** Pairwise proportion of testis TS-AS exons (a) or TS-GE genes (b) in the query species (x axis) that are conserved in the target species (y axis) across all species combinations. **c,d.** Number of testis TS-AS (c) and TS-GE (d) gains across ancestral nodes, with conservation classes represented by different shades. **e.** Number of exon (top) or gene (bottom) orthogroups containing at least one testis TS-AS exon (top) or TS-GE gene (bottom) in each branch, with conservation classes represented by different shades. **f.** Distributions of exon lengths across conservation classes of exon orthogroups containing at least one testis TS-AS exon in human (top) and fruitfly (bottom). The significance levels plotted on top reflect the results of a *Wilcoxon* test comparing the highly conserved groups to each of the others, and correspond to the following P-value cutoffs: **** = P-value ≤ 0.0001; *** = P-value ≤ 0.001; * = P-value ≤ 0.05; ns = non significant. The y axis has been truncated for visualization purposes. Exon orthogroups classified as “Others” were excluded from this plot due to their low numbers. **g.** Proportions of testis TS-AS exons (y axis) across species (x axis) that are conserved, convergent type I, convergent type II or new. In case of different classification of the same exons between species pairs, priority is given to the highest inferred conservation category. **h,i.** Number of testis TS-AS (h) or TS-GE (i) gains within gene orthogroups containing at least one testis TS-AS exon (h) or TS-GE gene (i). Gene orthogroups are classified as “recurrent” when undergoing more than one TS gain. **j.** Cumulative distributions of the proportions of gene orthogroups (y axis) containing testis TS-AS exons either in the observed TS set (blue) or in 10 randomized sets (gray) across species (x axis). The horizontal dashed lines represent the asymptotes for the observed TS and randomized sets as predicted by nonlinear least-squares fitting of an asymptotic exponential model, y=*a*(1−e^−*b*x^), where *a* is the asymptote and *b* the rate constant.

**Extended Data Fig. 7.**
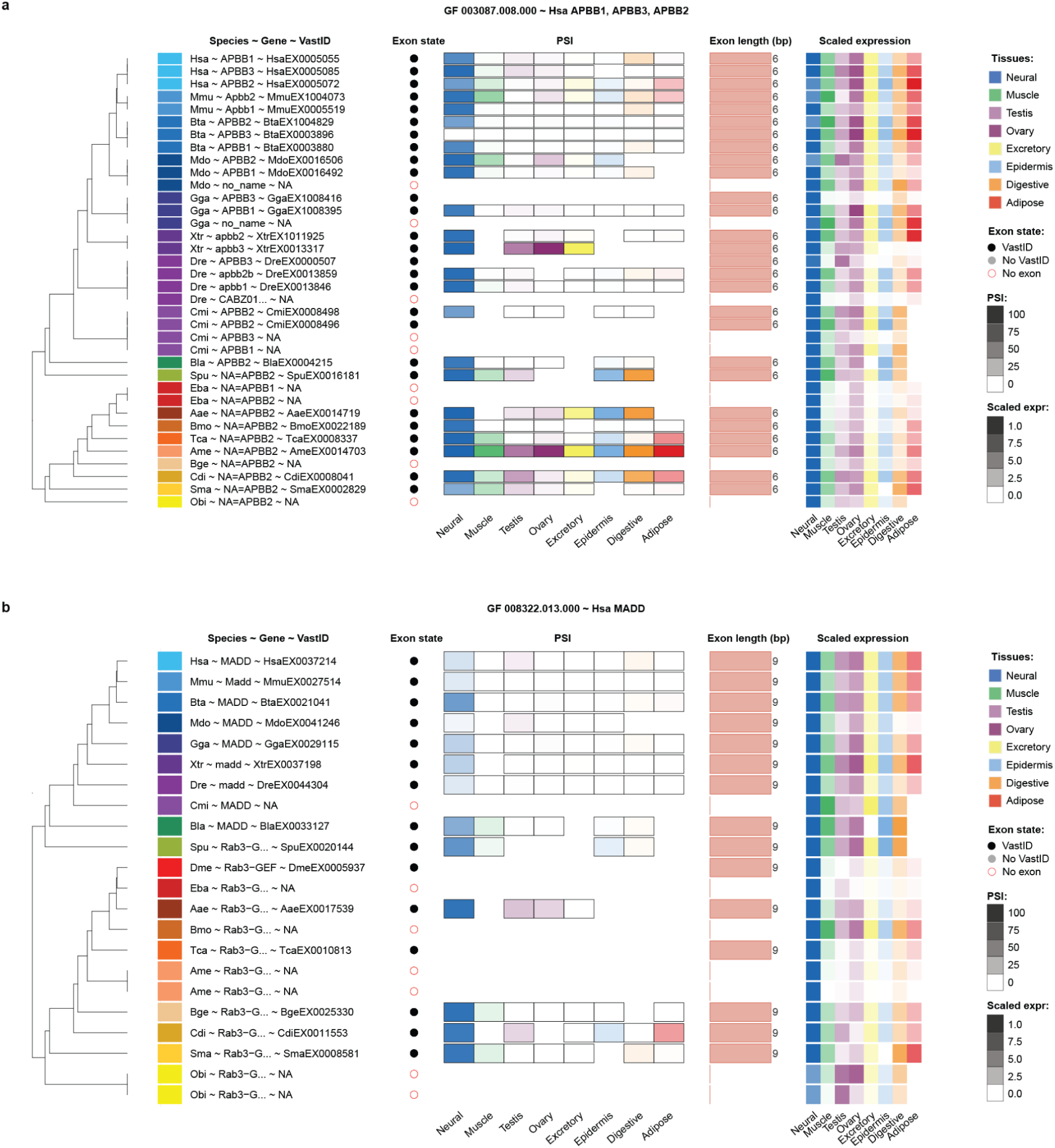
Summary plots of ancestral-bilaterian neural TS-AS exons: **a,b.** Summary plot of the ancestral-bilaterian microexons emerged in the APBB (a) and MADD (b) gene orthogroups. The “exon state” dotplot reports the annotation status of each given exon, where exons with “no VastID” are exons not included in the *vast-tools* ^23^ databases and for which AS quantification could not be carried out. Genes lacking the orthologous exons but still belonging to the same gene orthogroup were reported to provide a more complete evolutionary perspective. Scaled expression indicates expression proportions across tissues normalized to the maximum. Exons with membership score ≤ 0.15 in the *ExOrthist* re-clustered exon outgroups were excluded from the plots. The lack of black borders around the PSI cell of a given tissue indicates lack of minimum coverage (see **Methods**). The gene orthogroup ID is reported in the title.

**Extended Data Fig. 8.**
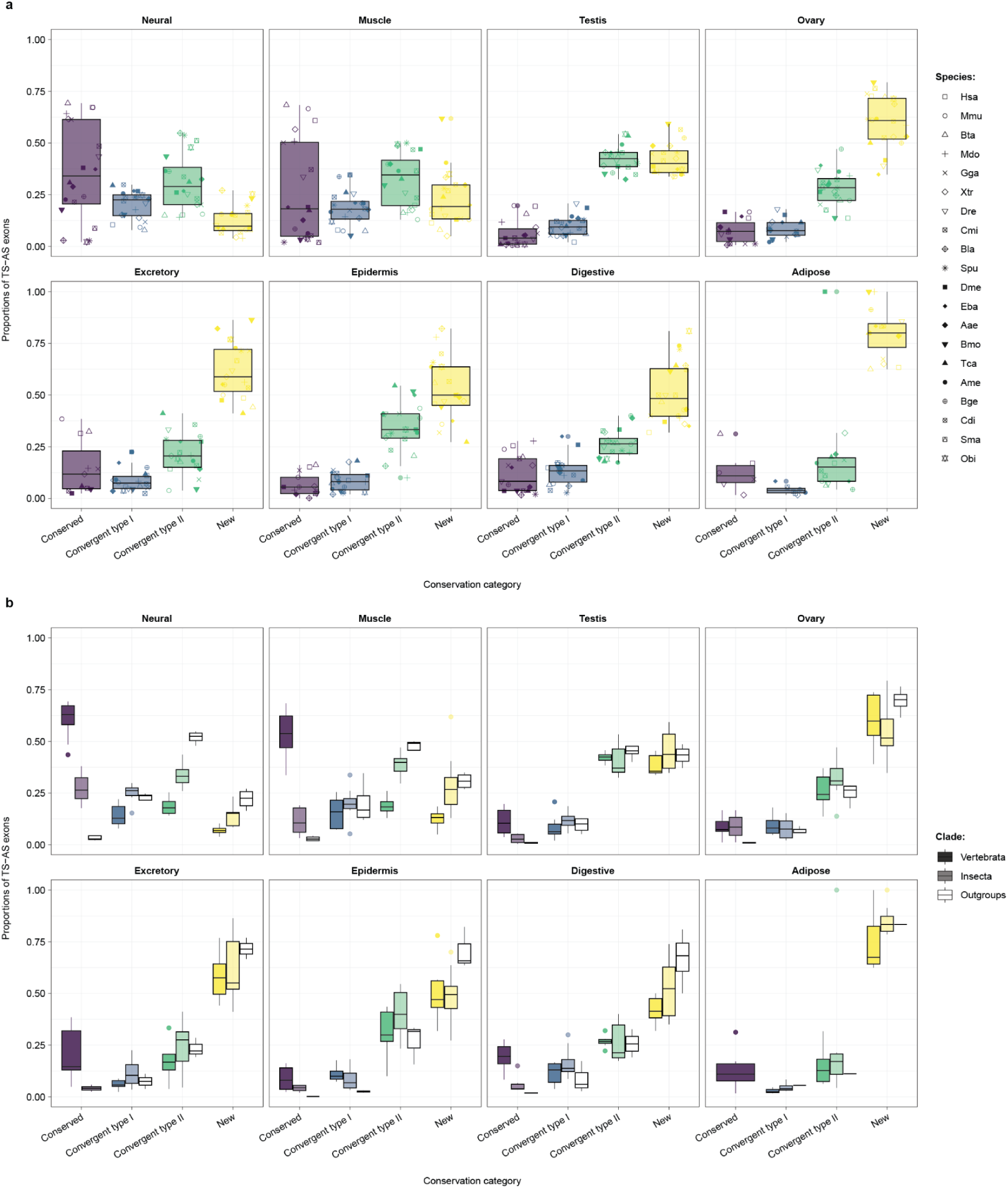
Relative proportions of TS-AS exons across regulatory conservation classes. **a,b.** Distributions of proportions of TS-AS exons across all species (a) or stratified by clade (b) across different conservation classes.

**Extended Data Fig. 9.**
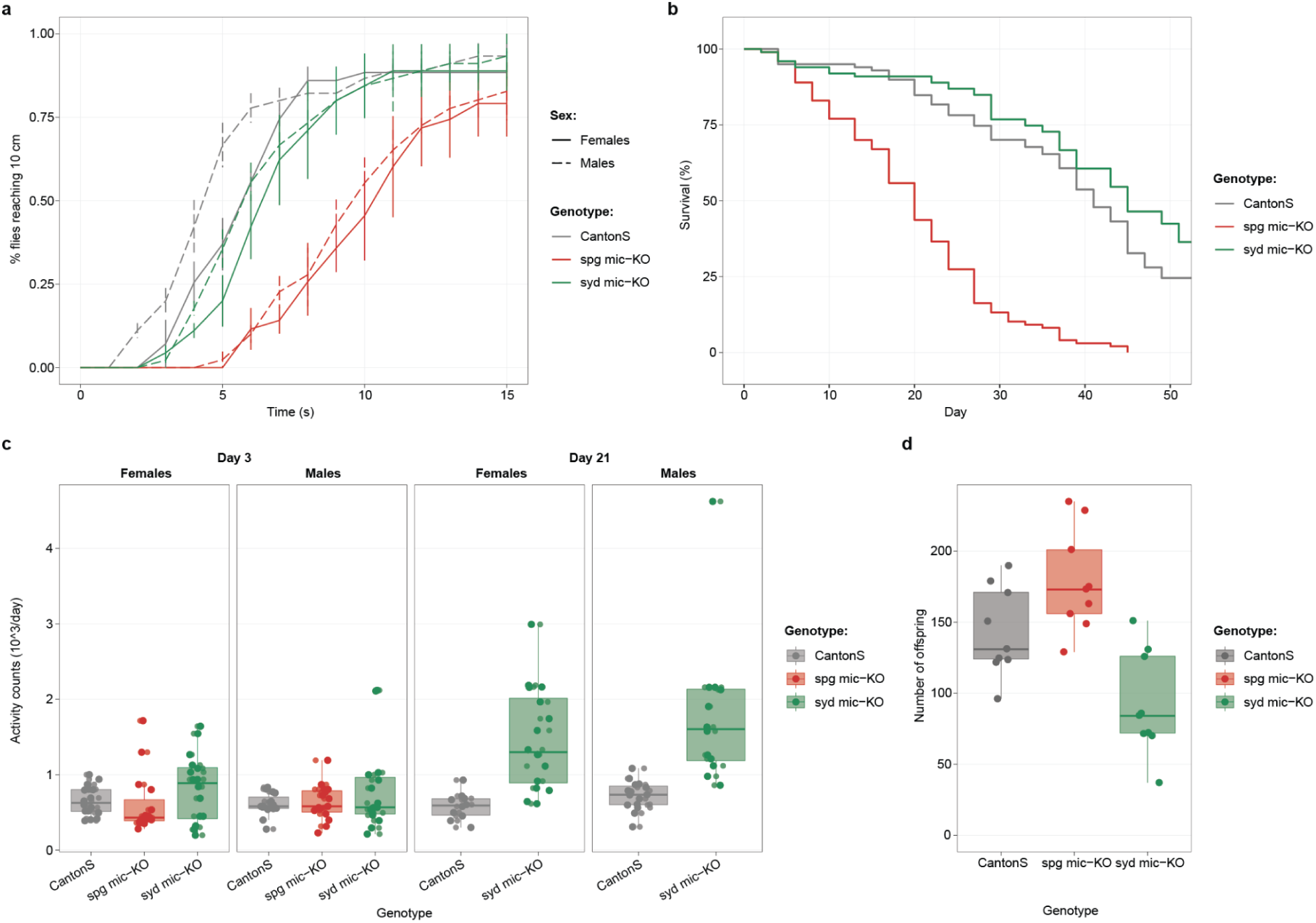
Phenotypic characterization of spg and syd microexon-KO flies. **a.** Anti-gravitaxis performance of 3-day-old CantonS control flies, spg microexon-KO flies and syd microexon-KO flies. Full and dashed lines respectively correspond to female and male flies. Points indicate the percentage of flies reaching 10 cm over time; error bars indicate the standard error of the mean. **b.** Survival curves for 100 CantonS control flies, 100 spg microexon-KO flies and 100 syd microexon-KO flies. **c**. Locomotor activity of CantonS control flies, spg microexon-KO flies and syd microexon-KO flies measured using a Drosophila Activity Monitor at days 3 and 21. Points represent individual flies. **d.** Number of offspring produced by age-matched female CantonS control flies, spg microexon-KO flies and syd microexon-KO flies. Points represent individual females.

## Supplementary Figures

**Supplementary Fig. 1.**
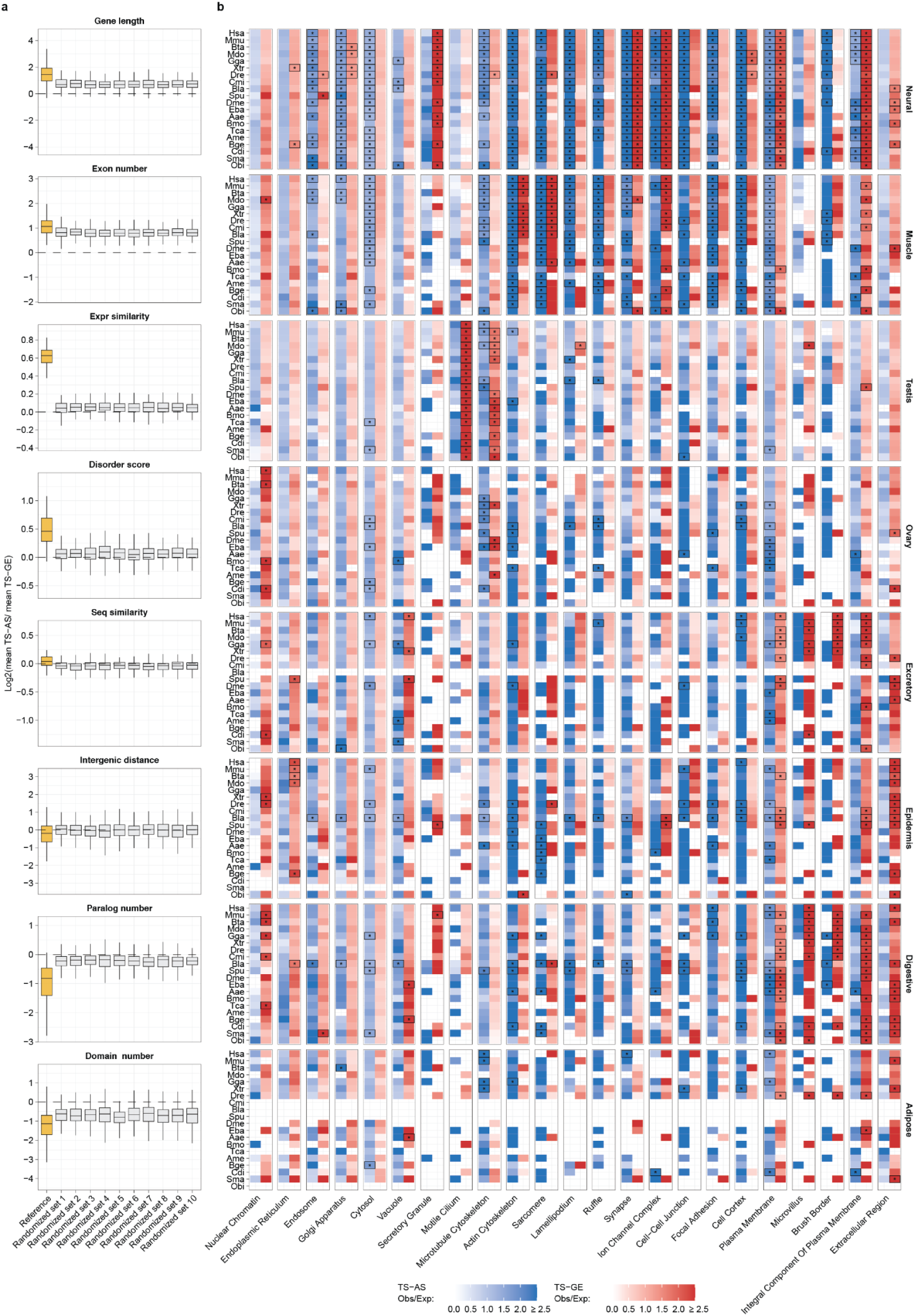
Extended gene features comparisons. **a.** Distributions of TS gene features (gold; see Fig. 2b) compared to the same distributions generated from ten randomized TS-AS and TS-GE gene sets. **b.** Observed vs Expected ratio of GO cellular components shown in Fig. 2d (x axis) across species (y axis) and tissues (vertical facet) for TS-AS (blue) and TS-GE (red) gene sets. Cells corresponding to significant GO enrichments (hypergeometric test, P-value ≤ 0.001) are marked by ticker borders and labelled with a star (*).

**Supplementary Fig. 2.**
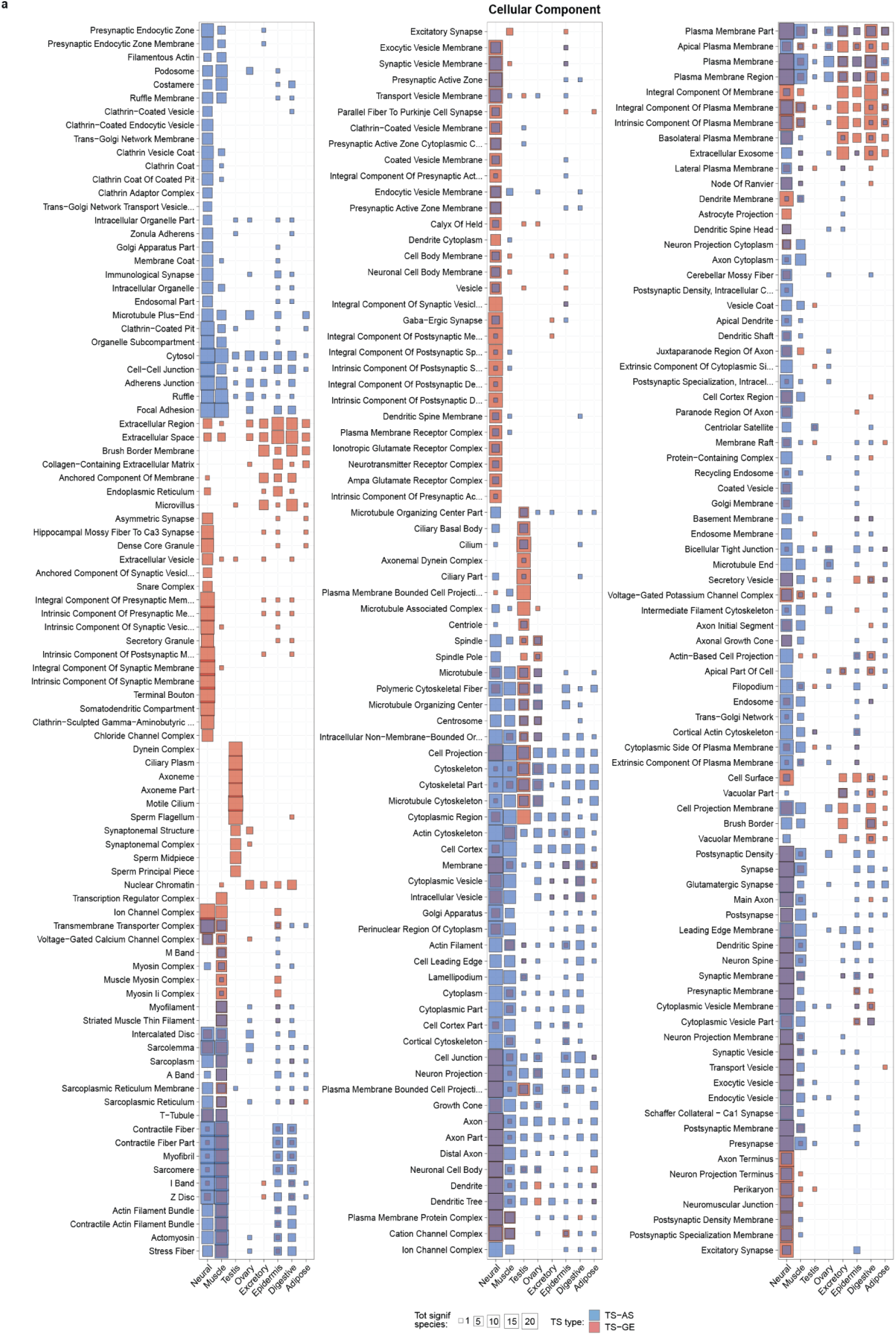
Cellular component GO enrichments of TS-AS and TS-GE genes. **a.** Number of species with significant enrichments for TS-AS (blue) and TS-GE (red) genes in each tissue (y axis) and across cellular component GO categories enriched in at least 5 species in any given tissue (x axis).

**Supplementary Fig. 3.**
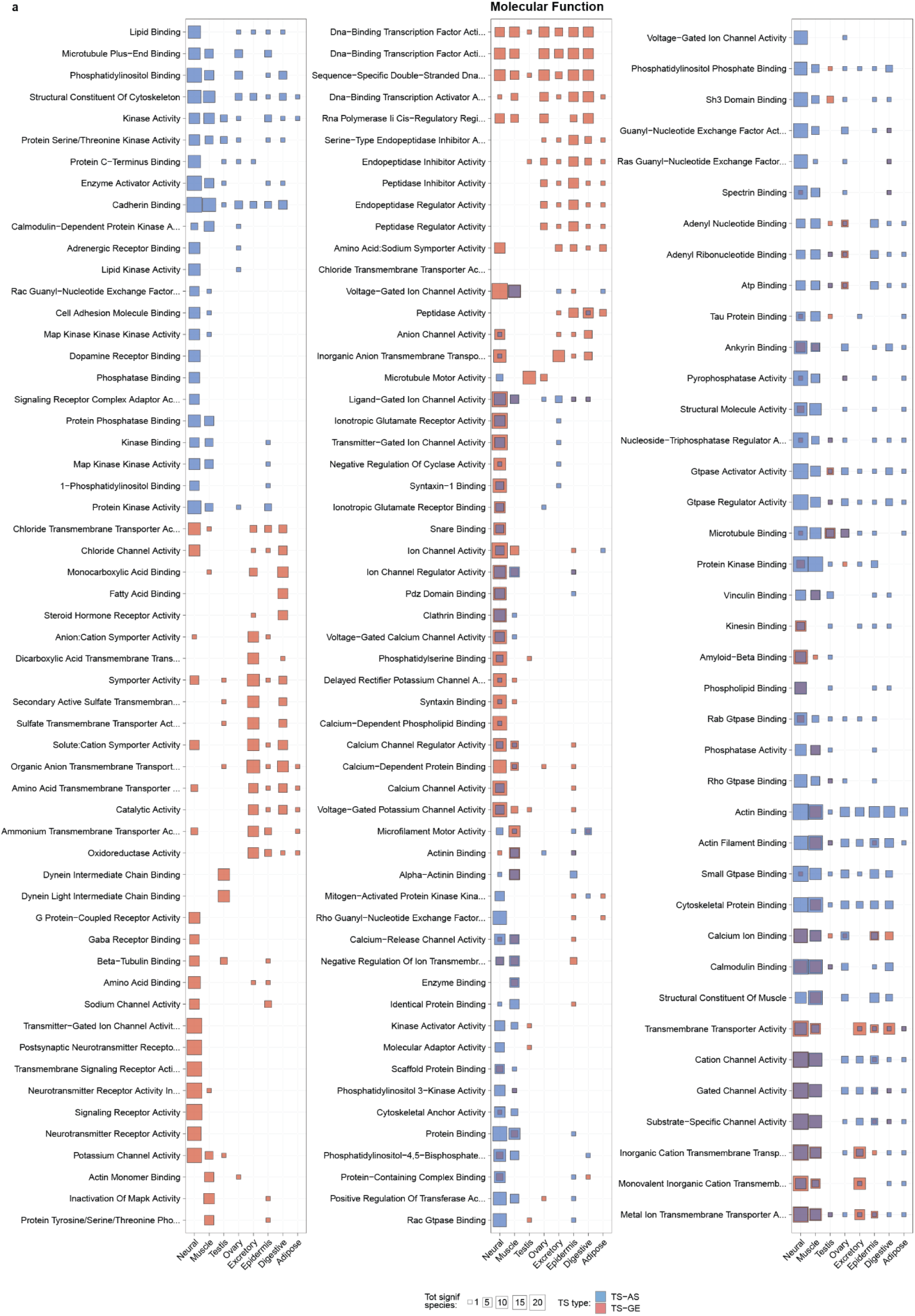
Molecular function GO enrichments of TS-AS and TS-GE genes. **a.** Number of species with significant enrichments for TS-AS (blue) and TS-GE (red) genes in each tissue (y axis) and across molecular function GO categories enriched in at least 5 species in any given tissue (x axis).

**Supplementary Fig. 4.**
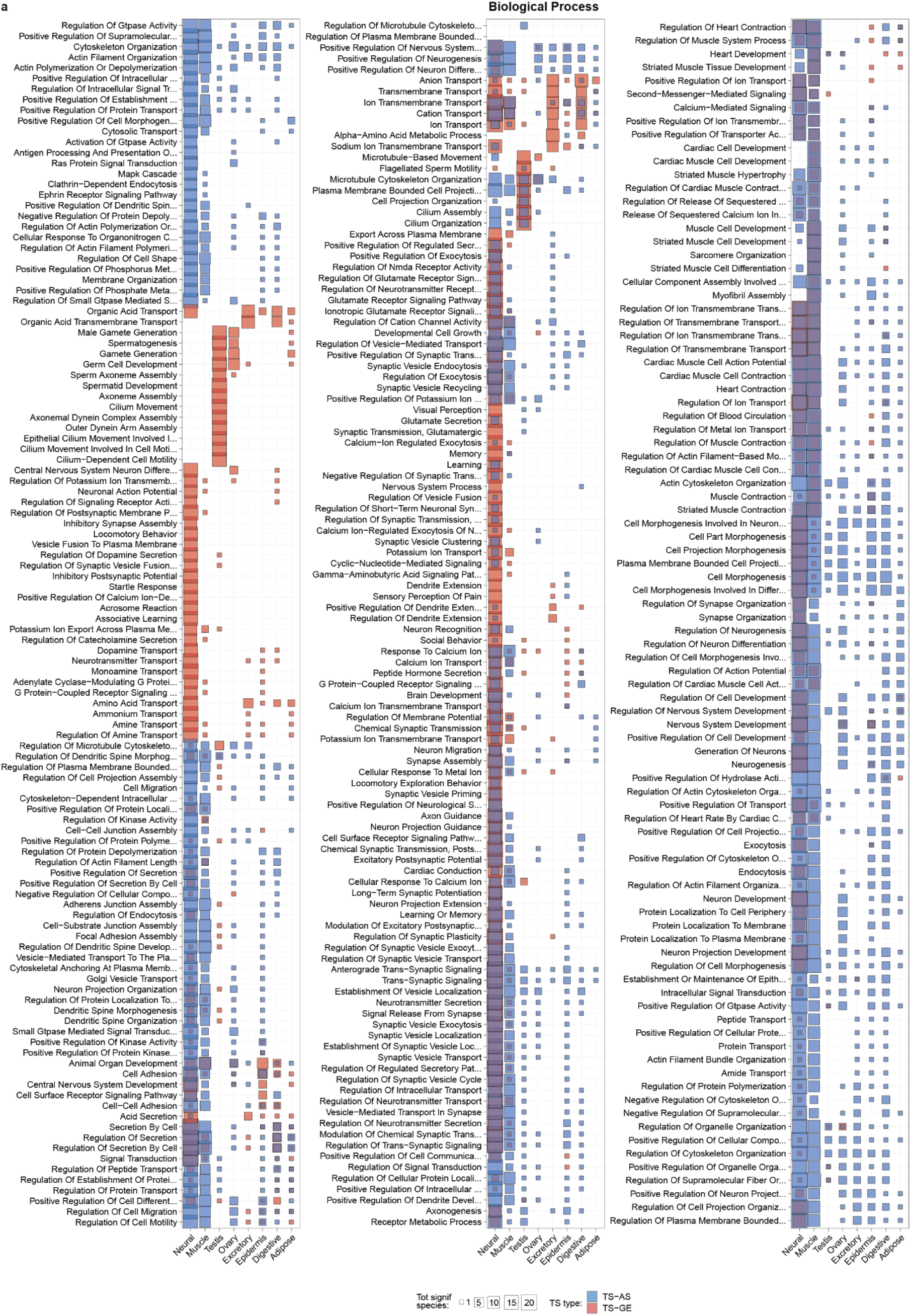
Biological processes GO enrichments of TS-AS and TS-GE genes. **a.** Number of species with significant enrichments for TS-AS (blue) and TS-GE (red) genes in each tissue (y axis) and across biological processes GO categories enriched in at least 12 species in any given tissue (x axis).

**Supplementary Fig. 5.**
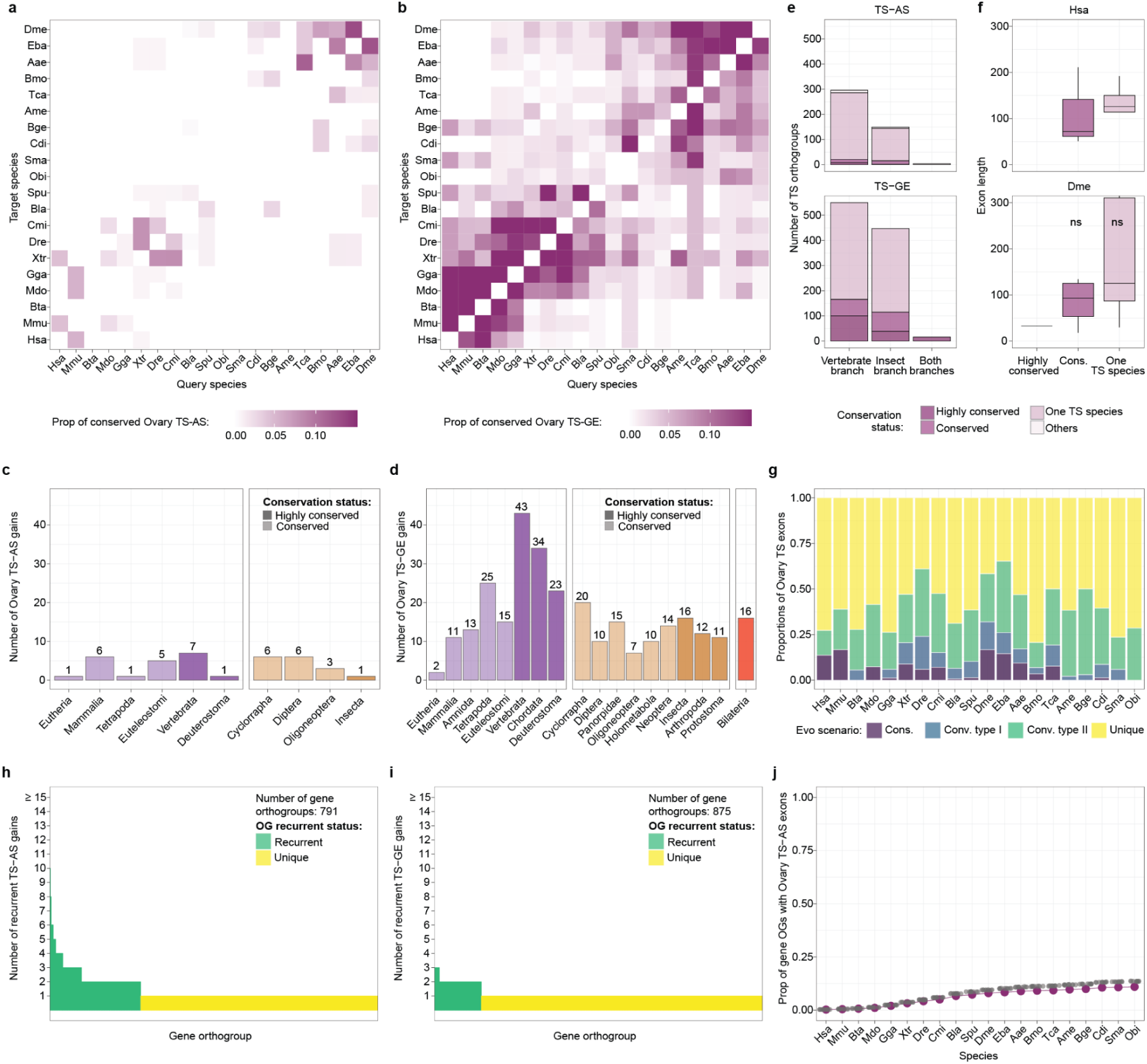
Evolutionary patterns of ovary TS-AS exons across bilaterians. **a,b.** Pairwise proportion of ovary TS-AS exons (a) or TS-GE genes (b) in the query species (x axis) that are conserved in the target species (y axis) across all species combinations. **c,d.** Number of ovary TS-AS (c) and TS-GE (d) gains across ancestral nodes, with conservation classes represented by different shades. **e.** Number of exon (top) or gene (bottom) orthogroups containing at least one ovary TS-AS exon (top) or TS-GE gene (bottom) in each branch, with conservation classes represented by different shades. **f.** Distributions of exon lengths across conservation classes of exon orthogroups containing at least one ovary TS-AS exon in human (top) and fruitfly (bottom). The significance levels plotted on top reflect the results of a *Wilcoxon* test comparing the highly conserved groups to each of the others, and correspond to the following P-value cutoffs: **** = P-value ≤ 0.0001; *** = P-value ≤ 0.001; * = P-value ≤ 0.05; ns = non significant. The y axis has been truncated for visualization purposes. Exon orthogroups classified as “Others” were excluded from this plot due to their low numbers. **g.** Proportions of ovary TS-AS exons (y axis) across species (x axis) that are conserved, convergent type I, convergent type II or new. In case of different classification of the same exons between species pairs, priority is given to the highest inferred conservation category. **h,i.** Number of ovary TS-AS (h) or TS-GE (i) gains within gene orthogroups containing at least one ovary TS-AS exon (h) or TS-GE gene (i). Gene orthogroups are classified as “recurrent” when undergoing more than one TS gain. **j.** Cumulative distributions of the proportions of gene orthogroups (y axis) containing ovary TS-AS exons either in the observed TS set (blue) or in 10 randomized sets (gray) across species (x axis).

**Supplementary Fig. 6.**
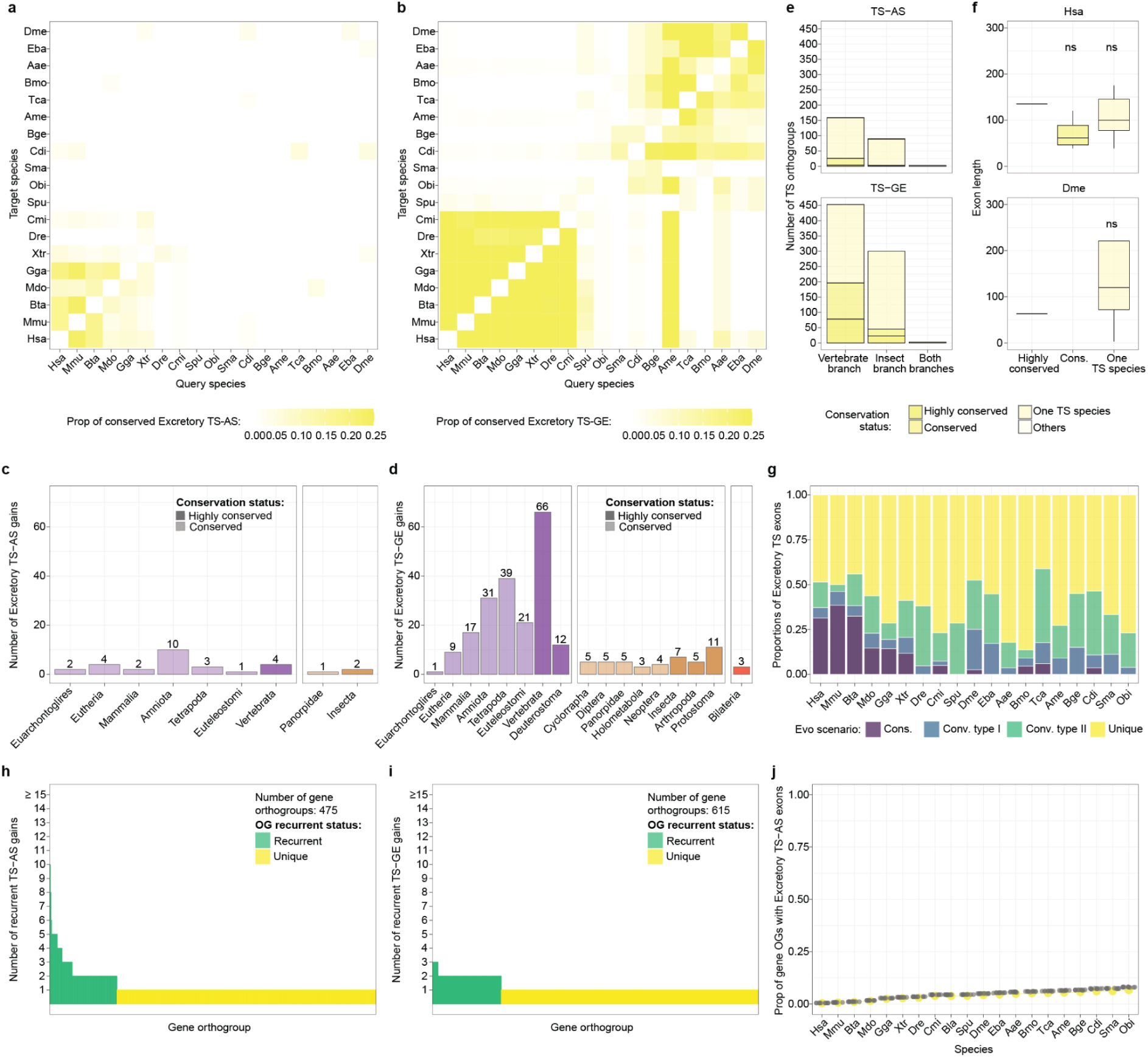
Evolutionary patterns of excretory TS-AS exons across bilaterians. **a,b.** Pairwise proportion of excretory TS-AS exons (a) or TS-GE genes (b) in the query species (x axis) that are conserved in the target species (y axis) across all species combinations. **c,d.** Number of excretory TS-AS (c) and TS-GE (d) gains across ancestral nodes, with conservation classes represented by different shades. **e.** Number of exon (top) or gene (bottom) orthogroups containing at least one excretory TS-AS exon (top) or TS-GE gene (bottom) in each branch, with conservation classes represented by different shades. **f.** Distributions of exon lengths across conservation classes of exon orthogroups containing at least one excretory TS-AS exon in human (top) and fruitfly (bottom). The significance levels plotted on top reflect the results of a *Wilcoxon* test comparing the highly conserved groups to each of the others, and correspond to the following P-value cutoffs: **** = P-value ≤ 0.0001; *** = P-value ≤ 0.001; * = P-value ≤ 0.05; ns = non significant. The y axis has been truncated for visualization purposes. Exon orthogroups classified as “Others” were excluded from this plot due to their low numbers. **g.** Proportions of excretory TS-AS exons (y axis) across species (x axis) that are conserved, convergent type I, convergent type II or new. In case of different classification of the same exons between species pairs, priority is given to the highest inferred conservation category. **h,i.** Number of excretory TS-AS (h) or TS-GE (i) gains within gene orthogroups containing at least one excretory TS-AS exon (h) or TS-GE gene (i). Gene orthogroups are classified as “recurrent” when undergoing more than one TS gain. **j.** Cumulative distributions of the proportions of gene orthogroups (y axis) containing excretory TS-AS exons either in the observed TS set (blue) or in 10 randomized sets (gray) across species (x axis).

**Supplementary Fig. 7.**
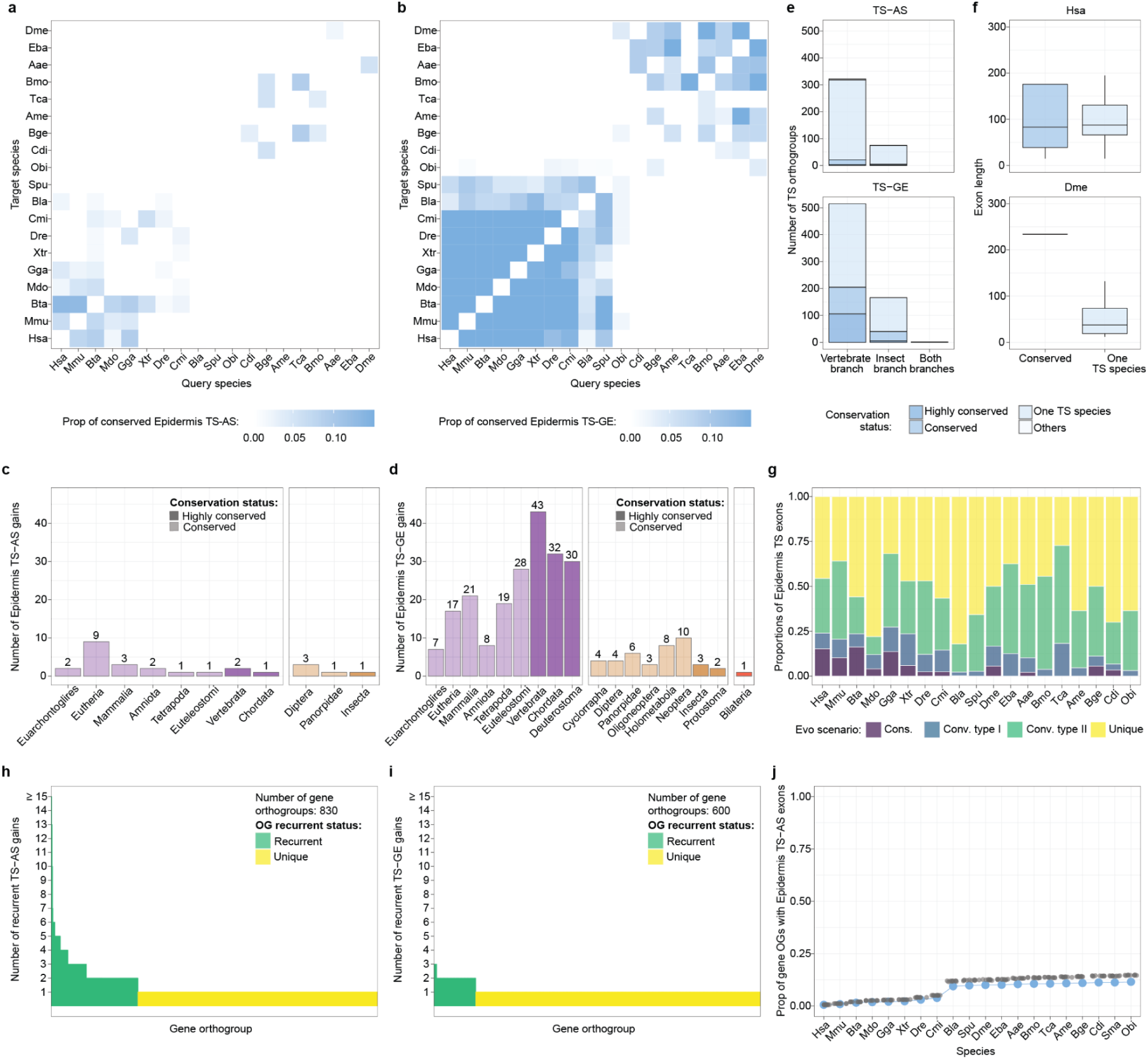
Evolutionary patterns of epidermis TS-AS exons across bilaterians. **a,b.** Pairwise proportion of epidermis TS-AS exons (a) or TS-GE genes (b) in the query species (x axis) that are conserved in the target species (y axis) across all species combinations. **c,d.** Number of epidermis TS-AS (c) and TS-GE (d) gains across ancestral nodes, with conservation classes represented by different shades. **e.** Number of exon (top) or gene (bottom) orthogroups containing at least one epidermis TS-AS exon (top) or TS-GE gene (bottom) in each branch, with conservation classes represented by different shades. **f.** Distributions of exon lengths across conservation classes of exon orthogroups containing at least one epidermis TS-AS exon in human (top) and fruitfly (bottom). The significance levels plotted on top reflect the results of a *Wilcoxon* test comparing the highly conserved groups to each of the others, and correspond to the following P-value cutoffs: **** = P-value ≤ 0.0001; *** = P-value ≤ 0.001; * = P-value ≤ 0.05; ns = non significant. The y axis has been truncated for visualization purposes. Exon orthogroups classified as “Others” were excluded from this plot due to their low numbers. **g.** Proportions of epidermis TS-AS exons (y axis) across species (x axis) that are conserved, convergent type I, convergent type II or new. In case of different classification of the same exons between species pairs, priority is given to the highest inferred conservation category. **h,i.** Number of epidermis TS-AS (h) or TS-GE (i) gains within gene orthogroups containing at least one epidermis TS-AS exon (h) or TS-GE gene (i). Gene orthogroups are classified as “recurrent” when undergoing more than one TS gain. **j.** Cumulative distributions of the proportions of gene orthogroups (y axis) containing epidermis TS-AS exons either in the observed TS set (blue) or in 10 randomized sets (gray) across species (x axis).

**Supplementary Fig. 8.**
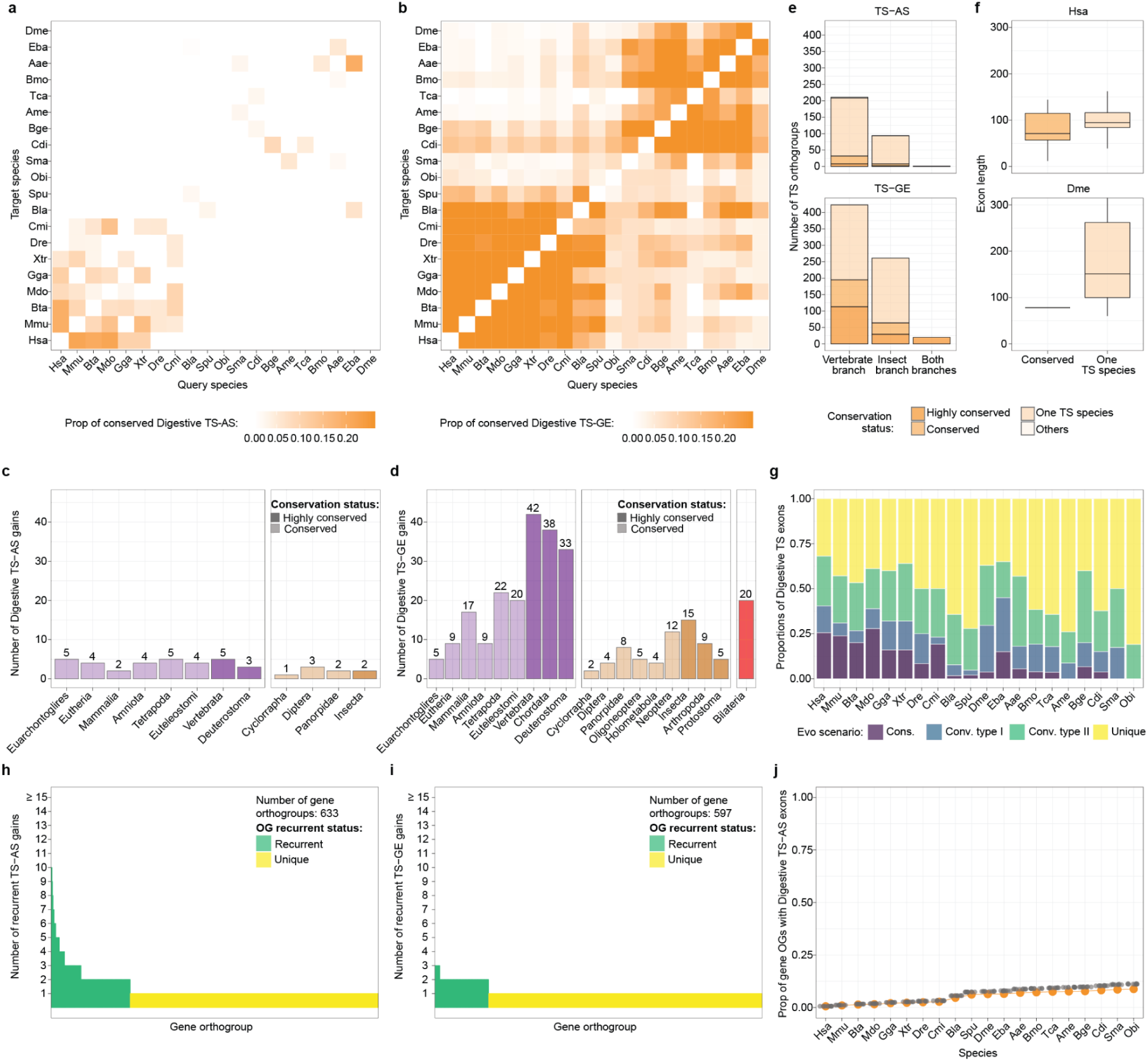
Evolutionary patterns of digestive TS-AS exons across bilaterians. **a,b.** Pairwise proportion of digestive TS-AS exons (a) or TS-GE genes (b) in the query species (x axis) that are conserved in the target species (y axis) across all species combinations. **c,d.** Number of digestive TS-AS (c) and TS-GE (d) gains across ancestral nodes, with conservation classes represented by different shades. **e.** Number of exon (top) or gene (bottom) orthogroups containing at least one digestive TS-AS exon (top) or TS-GE gene (bottom) in each branch, with conservation classes represented by different shades. **f.** Distributions of exon lengths across conservation classes of exon orthogroups containing at least one digestive TS-AS exon in human (top) and fruitfly (bottom). The significance levels plotted on top reflect the results of a *Wilcoxon* test comparing the highly conserved groups to each of the others, and correspond to the following P-value cutoffs: **** = P-value ≤ 0.0001; *** = P-value ≤ 0.001; * = P-value ≤ 0.05; ns = non significant. The y axis has been truncated for visualization purposes. Exon orthogroups classified as “Others” were excluded from this plot due to their low numbers. **g.** Proportions of digestive TS-AS exons (y axis) across species (x axis) that are conserved, convergent type I, convergent type II or new. In case of different classification of the same exons between species pairs, priority is given to the highest inferred conservation category. **h,i.** Number of digestive TS-AS (h) or TS-GE (i) gains within gene orthogroups containing at least one digestive TS-AS exon (h) or TS-GE gene (i). Gene orthogroups are classified as “recurrent” when undergoing more than one TS gain. **j.** Cumulative distributions of the proportions of gene orthogroups (y axis) containing digestive TS-AS exons either in the observed TS set (blue) or in 10 randomized sets (gray) across species (x axis).

**Supplementary Fig. 9.**
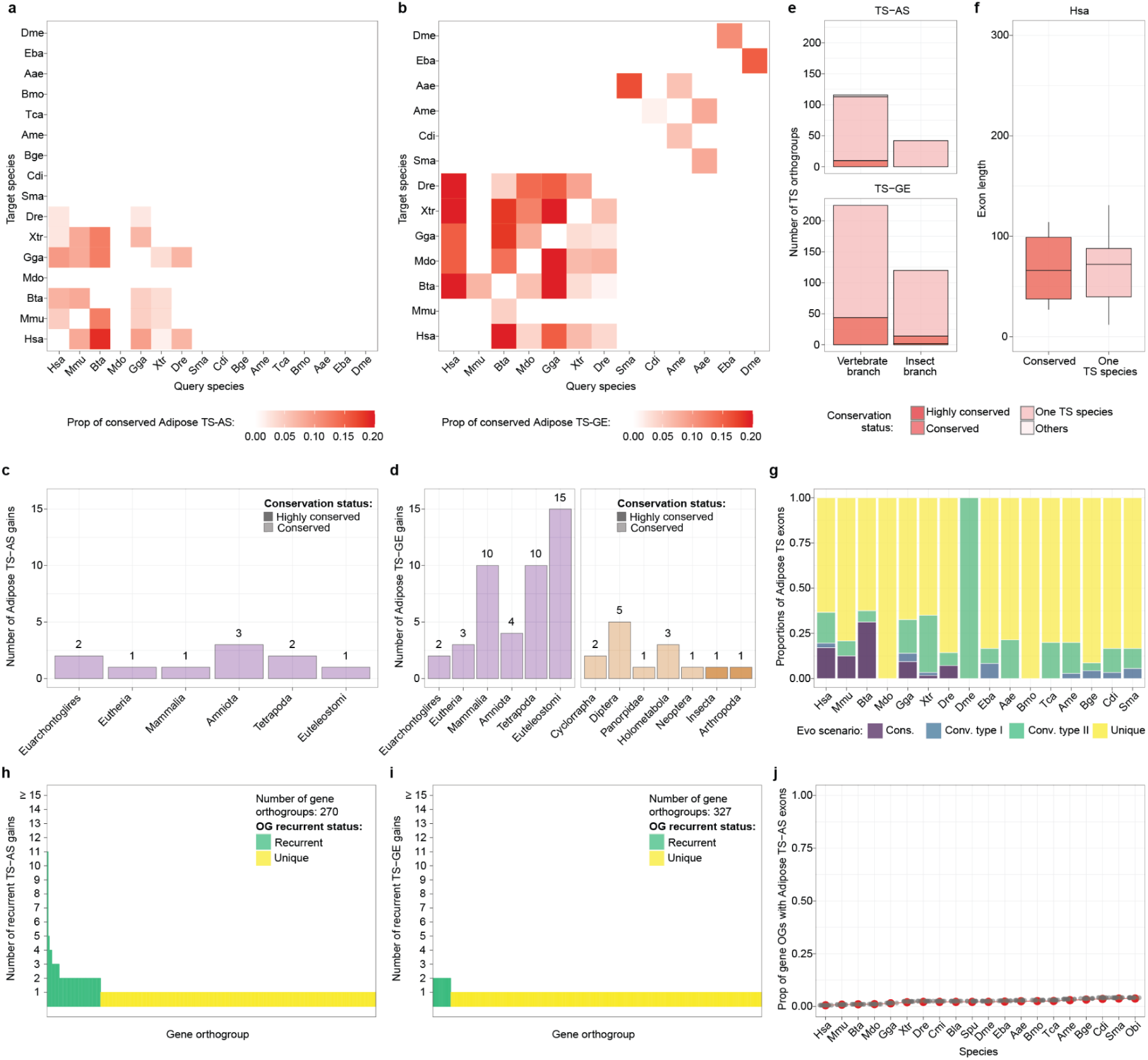
Evolutionary patterns of adipose TS-AS exons across bilaterians. **a,b.** Pairwise proportion of adipose TS-AS exons (a) or TS-GE genes (b) in the query species (x axis) that are conserved in the target species (y axis) across all species combinations. **c,d.** Number of adipose TS-AS (c) and TS-GE (d) gains across ancestral nodes, with conservation classes represented by different shades. **e.** Number of exon (top) or gene (bottom) orthogroups containing at least one adipose TS-AS exon (top) or TS-GE gene (bottom) in each branch, with conservation classes represented by different shades. **f.** Distributions of exon lengths across conservation classes of exon orthogroups containing at least one adipose TS-AS exon in human (top) and fruitfly (bottom). The significance levels plotted on top reflect the results of a *Wilcoxon* test comparing the highly conserved groups to each of the others, and correspond to the following P-value cutoffs: **** = P-value ≤ 0.0001; *** = P-value ≤ 0.001; * = P-value ≤ 0.05; ns = non significant. The y axis has been truncated for visualization purposes. Exon orthogroups classified as “Others” were excluded from this plot due to their low numbers. **g.** Proportions of adipose TS-AS exons (y axis) across species (x axis) that are conserved, convergent type I, convergent type II or new. In case of different classification of the same exons between species pairs, priority is given to the highest inferred conservation category. **h,i.** Number of adipose TS-AS (h) or TS-GE (i) gains within gene orthogroups containing at least one adipose TS-AS exon (h) or TS-GE gene (i). Gene orthogroups are classified as “recurrent” when undergoing more than one TS gain. **j.** Cumulative distributions of the proportions of gene orthogroups (y axis) containing adipose TS-AS exons either in the observed TS set (blue) or in 10 randomized sets (gray) across species (x axis).

**Supplementary Fig. 10.**
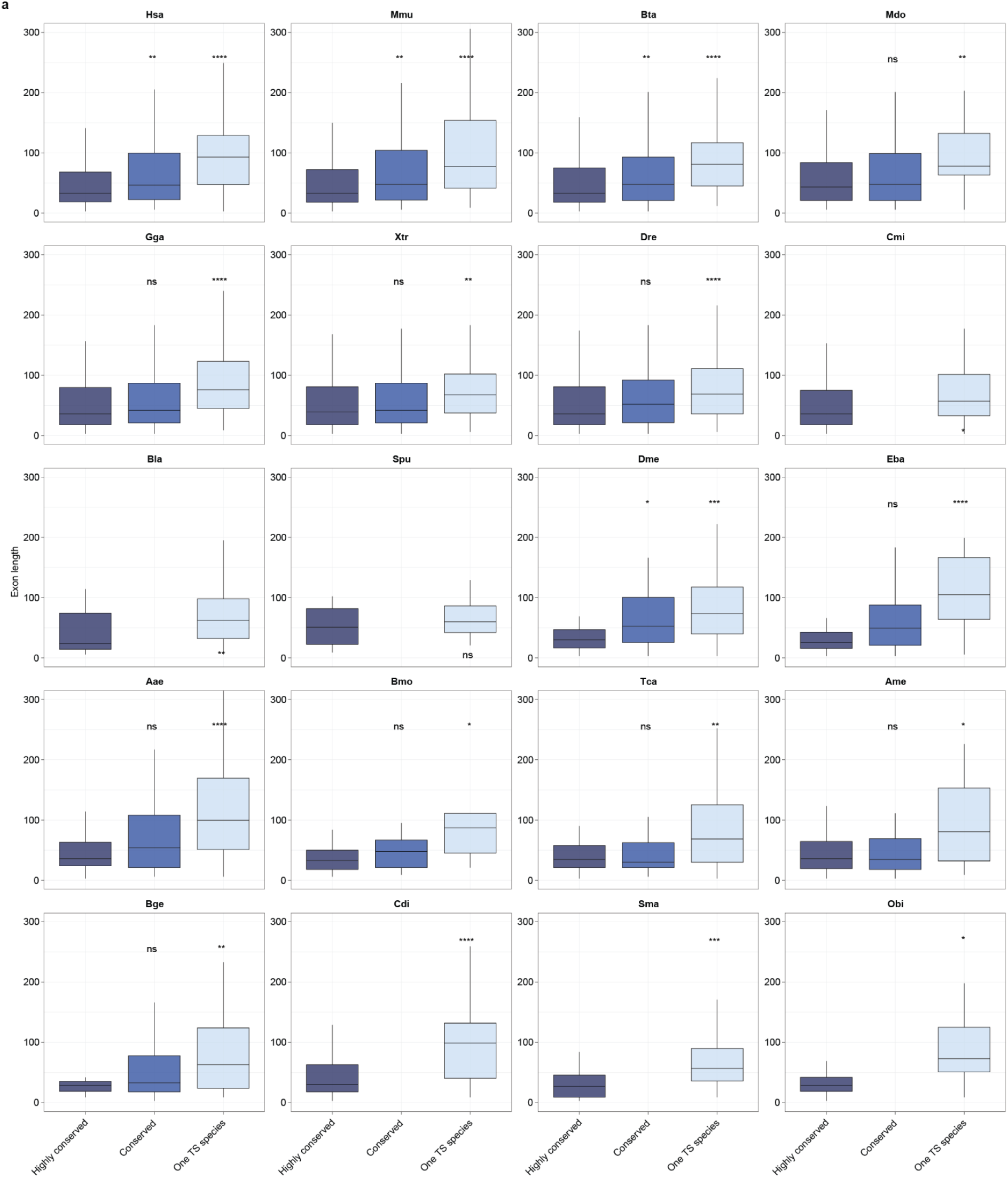
TS-AS neural exon lengths across conservation classes. Distributions of exon lengths in all species across conservation classes of exon orthogroups containing at least one neural TS-AS exon. The significance levels plotted on top reflect the results of a *Wilcoxon* test comparing the highly conserved groups to each of the others, and correspond to the following P-value cutoffs: **** = P-value ≤ 0.0001; *** = P-value ≤ 0.001; ** = P-value ≤ 0.01; * = P-value ≤ 0.05; ns = non significant.

## Supplementary Tables

**Supplementary Table 1**: Genome assembly for all the species.

**Supplementary Table 2**: Metadata for bulk RNA-seq dataset.

**Supplementary Table 3**: TS-AS and TS-GE genes across species and tissues.

**Supplementary Table 4**: GO enrichments across gene orthogroups classes (TS-AS only, TS-GE only, TS-AS and TS-GE, no TS).

**Supplementary Table 5**: Raw experimental data from fly microexon deletion experiments.

## Supplementary Datasets

**Supplementary Dataset 1:** Gene features across all species.

**Supplementary Dataset 2:** Exon orthogroups containing TS-AS exons.

**Supplementary Dataset 3:** Outputs of pairwise runs of *ExOrthist*’s *compare_exon_sets.pl*.

**Supplementary Dataset 4:** Inferred TS-AS gains for exon orthogroups across all tissues.

